# Loss of spontaneous vasomotion precedes impaired cerebrovascular reactivity and microbleeds in a mouse model of cerebral amyloid angiopathy

**DOI:** 10.1101/2024.04.26.591414

**Authors:** Mariel G Kozberg, Leon P Munting, Lee H Maresco, Corinne A Auger, Maarten L van den Berg, Baudouin Denis de Senneville, Lydiane Hirschler, Jan M Warnking, Emmanuel L Barbier, Christian T Farrar, Steven M Greenberg, Brian J Bacskai, Susanne J van Veluw

## Abstract

**Background:** Cerebral amyloid angiopathy (CAA) is a cerebral small vessel disease in which amyloid-β accumulates in vessel walls. CAA is a leading cause of symptomatic lobar intracerebral hemorrhage and an important contributor to age-related cognitive decline. Recent work has suggested that vascular dysfunction may precede symptomatic stages of CAA, and that spontaneous slow oscillations in arteriolar diameter (termed vasomotion), important for amyloid-β clearance, may be impaired in CAA.

**Methods:** To systematically study the progression of vascular dysfunction in CAA, we used the APP23 mouse model of amyloidosis, which is known to develop spontaneous cerebral microbleeds mimicking human CAA. Using *in vivo* 2-photon microscopy, we longitudinally imaged unanesthetized APP23 transgenic mice and wildtype littermates from 7 to 14 months of age, tracking amyloid-β accumulation and vasomotion in individual pial arterioles over time. MRI was used in separate groups of 12-, 18-, and 24-month-old APP23 transgenic mice and wildtype littermates to detect microbleeds and to assess cerebral blood flow and cerebrovascular reactivity with pseudo-continuous arterial spin labeling.

**Results:** We observed a significant decline in vasomotion with age in APP23 mice, while vasomotion remained unchanged in wildtype mice with age. This decline corresponded in timing to initial vascular amyloid-β deposition (∼8-10 months of age), although was more strongly correlated with age than with vascular amyloid-β burden in individual arterioles. Declines in vasomotion preceded the development of MRI-visible microbleeds and the loss of smooth muscle actin in arterioles, both of which were observed in APP23 mice by 18 months of age. Additionally, evoked cerebrovascular reactivity was intact in APP23 mice at 12 months of age, but significantly lower in APP23 mice by 24 months of age.

**Conclusions:** Our findings suggest that a decline in spontaneous vasomotion is an early, potentially pre-symptomatic, manifestation of CAA and vascular dysfunction, and a possible future treatment target.

## INTRODUCTION

Cerebral amyloid angiopathy (CAA) is a leading cause of intracerebral hemorrhage (ICH), a highly lethal type of stroke with a median one-month fatality rate of 40%^1^. In addition to symptomatic lobar ICH, CAA leads to smaller magnetic resonance imaging (MRI)-visible hemorrhagic lesions in the brain, including cerebral microbleeds and cortical superficial siderosis^2^. Pathologically, CAA is defined by an accumulation of amyloid-β in the walls of the brain’s vasculature, predominantly in cortical and leptomeningeal arterioles^3^.

CAA is associated with the dysfunction and loss of vascular smooth muscle cells (VSMCs)^3–5^, and patients with CAA have been observed to have a decline in stimulus-evoked cerebrovascular reactivity as measured by blood oxygen level dependent (BOLD) functional MRI studies^6^. Additionally, studies in pre-symptomatic individuals, defined as having no cognitive impairment or hemorrhagic lesions, with hereditary Dutch-type CAA have demonstrated declines in cerebrovascular reactivity ^7^, suggesting vascular dysfunction may precede symptomatic CAA.

In addition to their role in cerebrovascular reactivity, VSMCs also drive spontaneous slow oscillations in arterial diameter, termed vasomotion, at a frequency of ∼0.1 Hz^8,9^. Prior work has suggested that vasomotion plays a key role in glymphatic clearance^9,10^, with oscillations in vessel diameter increasing the mobility of the cerebrospinal fluid (CSF) located in the perivascular space, promoting exchange of molecules in the brain tissue and CSF^11^. Therefore, impairments in vasomotion could have implications for the accumulation of vascular amyloid-β in CAA^12^. Notably, amyloid-β levels in CSF are lower in CAA patients^13^, indicating that clearance of waste from the brain tissue towards the CSF may be impaired. In addition to vasomotion, beat-to-beat pulsatility, the change in vessel diameter with each heartbeat, has been suggested as another driving force for glymphatic clearance^14^. Beat-to-beat pulsatility has been shown to increase with age and cerebral small vessel disease, including CAA^15^.

We hypothesized that impairments in vasomotion would be observed with initial amyloid-β deposition, potentially contributing to a feed-forward mechanism in which reduced vasomotion leads to more deposition of amyloid-β, in turn leading to further declines in vasomotion. To study the progression of amyloid-β deposition, vascular dysfunction, and hemorrhage in CAA, we used the APP23 mouse model of amyloidosis, a model known to exhibit severe CAA and spontaneous MRI-visible cerebral microbleeds^16,17^. Longitudinal, unanesthetized *in vivo* 2-photon microscopy was performed in APP23 transgenic (Tg) mice and wildtype (WT) littermates from 7-14 months of age to track amyloid-β accumulation, arterial diameter, beat-to-beat pulsatility, and spontaneous vasomotion at an individual vessel level. We then performed *in vivo* MRI in separate cohorts of Tg mice and WT littermates at later ages (12, 18, and 24 months), assessing the timelines of the development of cerebral microbleeds using a multi-gradient echo (MGE) sequence, and assessing global cerebrovascular function using pseudo-Continuous Arterial Spin Labeling (pCASL)-MRI, combined with a hypercapnic (CO_2_) stimulus. Lastly, immunohistochemistry was performed to assess for vascular smooth muscle actin coverage at 12, 18, and 24 months in post-mortem brain tissue of Tg mice and WT littermates.

## METHODS

### Animals

All experiments were approved by the animal care and use committee of the Massachusetts General Hospital under protocol number 2018N000131. The study was compliant with the National Institutes of Health Guide for the Care and Use of Laboratory Animals and reported according to the ARRIVE guidelines^18^. Some of the mice were acquired directly from the Jackson Laboratory, Bar Harbor, Maine (strain #030504), others were derived from in-house breeding (also founded with strain #030504 mice from the Jackson Laboratory). See further mouse details including housing information and which mice were bred in house in the Online Supplement - Detailed Methods section. Different cohorts of mice were used for 2-photon microscopy and MRI.

#### 2-photon microscopy

Mice were imaged monthly longitudinally from ∼7-14 months of age as follows: 4 WT mice (1 female) and 4 Tg mice (2 female). The average age at first imaging session was 250 ± 30 days and 252 ± 29 days in WT and Tg mice respectively. An additional cohort was imaged at 15-16 months of age: 4 WT (1 female), 3 Tg (2 female). A third cohort of 15-16-month-old mice was used for experiments assessing the effects of isoflurane on vascular diameter: 3 WT (1 female), 3 Tg (1 female).

#### MRI

A cross-sectional study design was used with three age groups of mice as follows: 1) 12 months: 6 WT mice (4 female), 5 Tg mice (4 female); 2) 18 months: 10 WT mice (5 female), 10 Tg mice (4 female); and 3) 24 months: 9 WT littermates (3 females), 6 Tg (4 female).

Five mice (1 WT 12-month-old male, 1 WT 18-month-old male, 1 WT 18-month-old female, 2 Tg 18-month-old males) in the MRI cohort were excluded from the pCASL analysis, because of technical problems (see paragraph on MR image processing). One additional 18-month-old Tg female was excluded from the MRI cohort, due to detection of a large brain tumor-like lesion with MRI.

#### Immunohistochemistry

For histopathological analysis, the following mice were included: 1) 12 months: 3 WT (2 female), 3 Tg (3 female); 2) 18 months: 7 WT (3 female), 6 Tg (3 female); and 3) 24 months: 8 WT (3 female), 5 Tg (3 female).

### 2-photon microscopy

#### Surgical preparation

Cranial windows and headposts were implanted in mice as previously described^9^. Briefly, mice were anesthetized with 1.5-2% isoflurane (in 100% O_2_) throughout surgical procedures. A dental drill was used to drill a 6 mm craniotomy overlying the bilateral somatosensory cortices, which was then replaced with an 8 mm glass coverslip, fixed in place with a mixture of dental cement and Krazy Glue. Dura was kept intact. Custom-designed stainless-steel headposts were fixed to the skull using C&B Metabond (Parkell). Buprenorphine (0.05 mg/kg) and Tylenol (300 mg/mL) were used for post-surgery analgesia.

#### Image acquisition

At least 4 weeks following each surgery, longitudinal 2-photon imaging was initiated. Prior to imaging, mice were habituated to awake imaging through at least two 5-15 min sessions of head fixation via headpost to a stereotactic frame on a circular treadmill. Mice were then imaged approximately monthly for up to 6 months. One day prior to each imaging session, 300 μL methoxy-XO4 solution (Glixx lab) (∼ 5 mg/kg, dissolved in 3% Cremophor®EL (Sigma) and phosphate-buffered saline) was injected intraperitoneally to fluorescently label amyloid-β. On the day of imaging, mice were induced with 5% isoflurane and retro-orbital intravenous injections of 150 μL of fluorescein or Texas Red dextran (70 kDa) solution (12.5 mg/mL; Invitrogen) were performed. Mice were then allowed to wake-up over a 15-minute period prior to imaging. During each imaging session, mice were fixed via headposts in a stereotactic frame and allowed to move freely on a circular treadmill.

Mice were imaged using a FluoView FV1000MPE 2-photon laser-scanning system (Olympus) mounted on a BX61QI microscope (Olympus), equipped with a 25x (numerical aperture = 1.05) dipping water immersion objective (Olympus). A MaiTai® DeepSee™ Ti:Sapphire mode-locked laser (Spectra-Physics) generated 2-photon excitation at 800 nm. External detectors containing three photomultiplier tubes (PMTs, Hamamatsu) collected emitted light at 420-460 nm (blue), 495-540 nm (green), and 575-630 nm (red). Each field of view (FOV) was imaged ∼monthly, unless precluded by clouding of the window. See imaging parameters in the Online Supplement – Detailed Methods section. Longitudinal two-photon imaging of mice concluded after 6 months of imaging, if clouding of the window precluded further imaging (n = 2, each after 3 months of imaging), or with death of the mouse (n = 1, after 4 months of imaging).

#### Image processing

2-photon images were analyzed in FIJI and MATLAB using in-house developed scripts. To assess spontaneous vasomotion, vascular diameter was assessed in arteriolar segments in the 5 min timeseries data using the dextran channel data (either green or red channel); the methoxy-XO4 channel (blue) was subtracted because of bleed-through of the methoxy-XO4 signal in the other channels. Fast-Fourier transforms were performed and the peak amplitude in the frequency domain between 0.04 and 0.13 Hz was calculated for each segment. An average “vasomotion peak” was calculated for each FOV arteriole (∼ 2-4 segments/arteriole). To assess % CAA (i.e. vascular amyloid-β) coverage of pial arterioles, maximum intensity projections including the pial vasculature in each FOV were used. Regions of interest (ROIs) were drawn on the dextran channel (green or red channel) surrounding arterioles (excluding venules), and the total # of dextran-positive pixels in a binarized image (corresponding to overall arteriole area) was calculated. Then, these ROIs were applied to the methoxy-XO4 channel (blue) and the total # of CAA-positive pixels was calculated in a binarized image, to obtain a CAA coverage %. Baseline arteriolar diameters were calculated from line scan assessments perpendicular to the vessel segment. Red blood cell (RBC) velocities were calculated from line scans acquired parallel to the vessel wall. Beat-to-beat pulsatility was measured as 1) the absolute value of the area under the curve of Δ diameter vs. time plots and 2) the amplitude of the peak corresponding to heart rate in the frequency domain (calculated through a fast-Fourier transform).

### MRI

#### Image acquisition

For MRI acquisition, the mice were anesthetized with a low isoflurane protocol previously described in detail^19^. In short, anesthesia was induced with 2% isoflurane and maintained at 1.1% in oxygen-enriched air (30% O_2_). The mice were head-fixed and freely breathing. Breathing rates were monitored using a pressure-sensitive pad placed below the animal. Temperature was maintained at 36.5°C using a feedback-controlled air heat pump (Kent Scientific). A 9.4 T MRI scanner with Paravision 6.0.1 software (Bruker, Ettlingen, Germany) was used with a head phased array receive coil and whole-body volume transmit coil. See scan details in the detailed methods section.

In a subset of 6 mice from the MRI cohort (3 WT and 3 Tg 18-month-old mice), 1 week before the MRI scan, mice were placed in the same animal cradle as used during MRI, and underwent the same anesthesia protocol as described above, including 10% CO_2_ stimulation, but they did not undergo MRI scanning. Instead, the right dorsal flank was shaved, and transcutaneous (tc)-pCO_2_ was measured using a skin probe coupled to a blood gas analyzer (CombiM54, Radiometer America, CA).

#### Image processing

The pCASL image analysis pipeline has been described in detail previously^19^. In short, consecutive EPIs from individual scans were first spatially aligned. CBF maps were generated for each label and control pair, using MATLAB-based open-source software^20^. CBF was expressed in mL/100 g/min, derived from the label and control signal difference using Buxton’s perfusion model^21^ as follows:

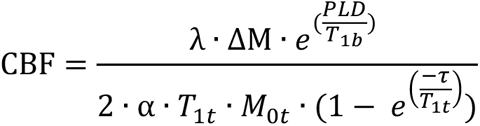

Here, λ is the blood–brain partition coefficient, i.e. 0.9 mL/g^22^, ΔM is the pCASL label and control signal difference and T_1b_ is the longitudinal relaxation time of blood, taken to be 2,430 ms at 9.4 T^23^. The labeling efficiency (α) was assumed to be 0.80, which was the median α of all WT animals (Supplemental Figure 6) and is in correspondence with literature^24^. ROIs were drawn by hand on the reference T_2_-weighted RARE scan to extract regional CBF values. CBF time-profiles were smoothed by removing outliers (exclusion of values higher or lower than two times the standard deviation) and applying a three-point moving average. CVR was derived by normalizing the CBF values to the average CBF between minute 1-5. See further image processing details in the Online Supplement – Detailed Methods section.

Five mice were excluded from the pCASL analysis, because of ghosting artefacts in the EPIs. Because the ghosts were variable in signal intensity, the brain signal varied randomly over time, resulting in large subtraction errors when deriving the perfusion signal. The ghosts likely appeared in these five datasets due to insufficient head fixation, resulting in respiration-related movement.

TOF images were filtered to enhance curvilinear structures, as in ^25^, using ImageJ’s “Tubeness” filter. Subsequently, maximum intensity projections were created and qualitatively assessed to screen for abnormalities. The MGE scans were inspected in MATLAB by 2 blinded observers (M.G.K. and M.L.B) to determine the number of microbleeds, defined as hypointense round or ovoid lesions. The intraclass correlation coefficient (ICC) was excellent (0.833). Final microbleed counts were determined at a consensus meeting.

### Immunohistochemistry

After completion of MRI scans, mice were euthanized on the same day through CO_2_ asphyxiation, after which they were transcardially perfused with 20 mL phosphate-buffered saline (PBS). When possible, 2-photon imaged mice were also euthanized and transcardially within 1 week of the last 2-photon imaging session. For both MRI and 2-photon imaged mice, the brain was extracted and fixed in 4% paraformaldehyde with 15% glycerol in PBS for several months, then processed, and embedded in paraffin in coronal orientation. Coronal serial sections of 6 μm thickness were cut with a microtome.

Serial sections from a subset of mice (described above in the *Animals* section) underwent immunohistochemistry against amyloid-β and smooth muscle actin (SMA). See staining details in the Online Supplement – Detailed Methods section. Sections were imaged using the Hamamatsu NanoZoomer Digital Pathology (NDP)-HT scanner (C9600–12, Hamamatsu Photonics K.K., Japan) at 20x magnification. The viewing platform NDP.View (version 2.6.13) was used to analyze the digital sections. A trained rater (C.A.A.) selected all pial arterioles from one amyloid-β stained coronal section at approximately the level of the somatosensory cortex from each mouse. Vessels were rated on the Vonsattel vessel grading scale^3^ as follows: Vonsattel grade 0 (no amyloid-β), Vonsattel grade 1 (patchy amyloid-β deposition), and Vonsattel grade 2 vessels (circumferential amyloid-β). Then, at least 1 week after vessel selection, blinded to the mouse age, genotype, and Vonsattel scores, the rater scored each selected vessel on adjacent SMA stains on the following SMA scale (modified from ^5^): 0 (no SMA), 1 (incomplete SMA coverage), 2 (complete SMA coverage). Arterioles in 2 randomly selected amyloid-β stains (both Tg) and 2 SMA stains were rated by a second rater (M.G.K.) for both Vonsattel grades and SMA scores. Inter-rater reliability was excellent for both Vonsattel grades (kappa = 0.96) and SMA scores (kappa = 0.87). An average SMA score per Vonsattel grade was then generated for each mouse.

### Statistics

Statistical analyses were performed in R, Prism v10, and MATLAB. Linear mixed effects (LME) models were fitted using the “lme4” package version 1.1-26 in R to assess changes in vascular amyloid-β, vessel diameter, RBC velocity, vasomotion, and beat-to-beat pulsatility. All models tested for heteroskedasticity, and log transforms were used as needed. In all models, subject (i.e. mouse ID) and arteriole were treated as random effects. Fixed effects used in each model are detailed in the results section, Table 1, and Supplemental Tables 1-6. Model fits were compared through evaluating the Akaike information criterion (AIC) and Bayesian information criterion (BIC) for each model; lower values indicated a better fit.

**Table 1.**
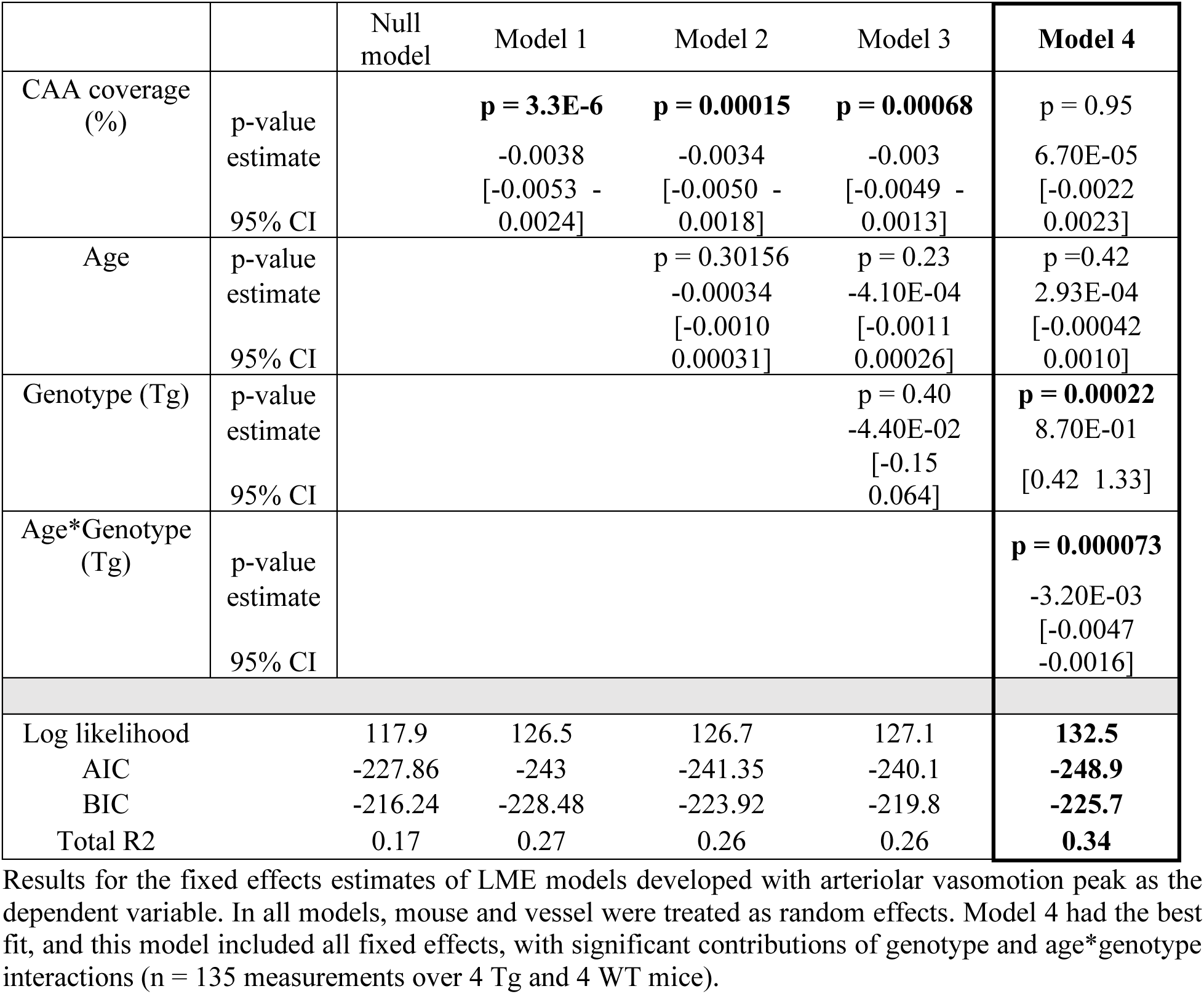
Vasomotion models.

Further details regarding statistical analyses are included in the Online Supplement – Detailed Methods section.

## RESULTS

### Vascular amyloid-β starts to accumulate in pial arterioles at ∼ 8-10 months of age in APP23 Tg mice

CAA was first observed in pial arterioles in Tg mice between ∼ 8 – 10 months of age in 3 of the 4 Tg mice imaged (Figure 1). To assess the relationship between age and vascular amyloid-β (i.e. CAA) coverage, an LME model was applied with age as the predictor variable, mouse and vessel as random effects, and % CAA coverage as the dependent variable. % CAA coverage increased significantly with age (n = 62 measurements across 4 Tg mice, p = 9.7E-16) (Figure 1B). The model including age explained 80% of the variance in % CAA coverage (comparison between null and full model: c^2^ = 65.7, p = 5.2E-16). See Supplemental Table 1 for additional model details. Note, one mouse did not develop CAA within the captured FOVs over the course of imaging. This mouse was re-genotyped confirming it was a Tg mouse and therefore this mouse was included in all analyses presented.

**Figure 1.**
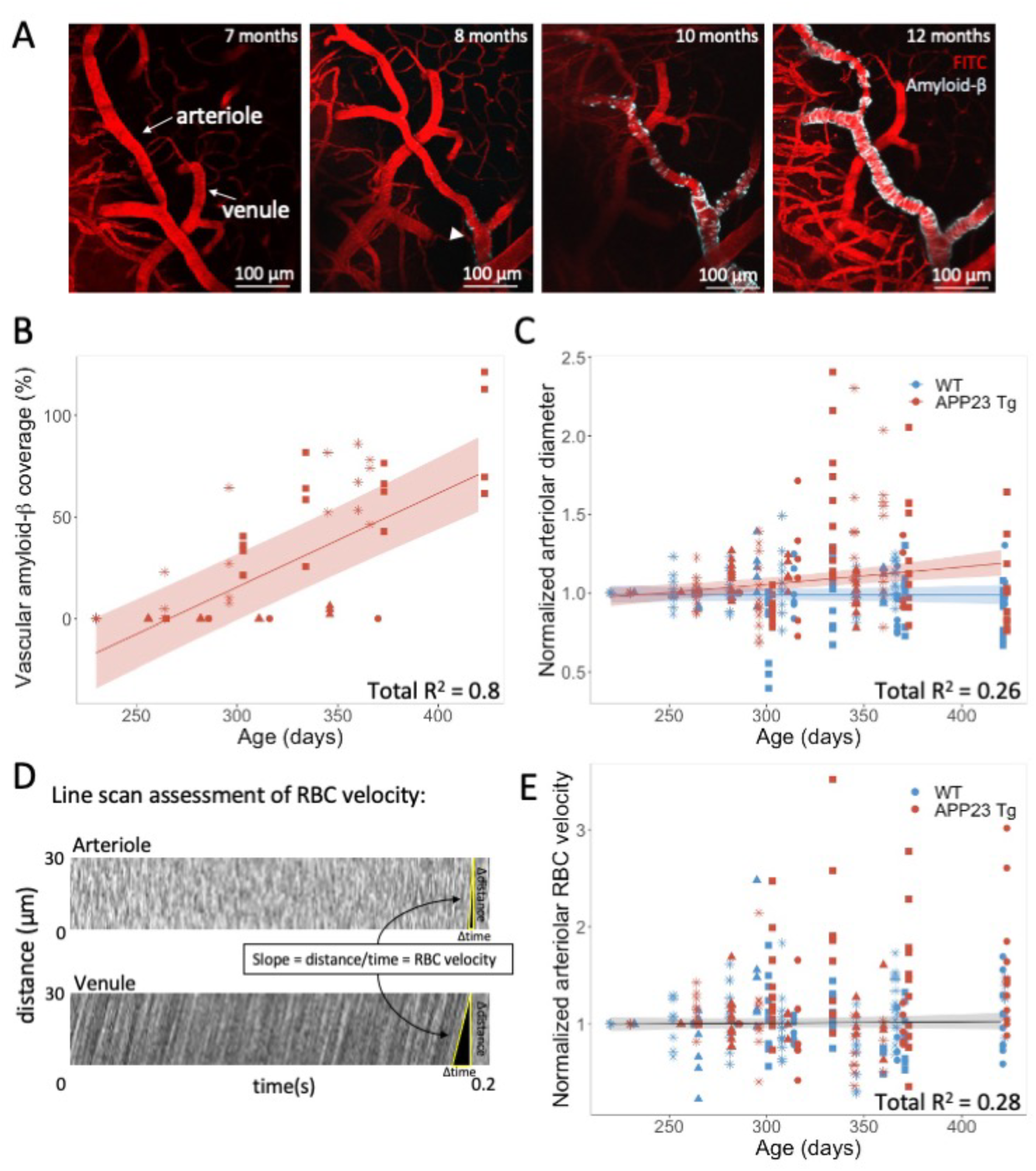
Vascular amyloid-β deposition starts to accumulate in APP23 Tg mice around 8-10 months of age and increasing age in APP23 Tg mice is associated with an increase in baseline vessel diameter. A) *In vivo* 2-photon images of the same arteriole and venule in a representative APP23 Tg mouse imaged from 7 to 12 months of age. Amyloid-β was first observed along the arteriole (arrowhead) at 8 months. B) % vascular amyloid-β coverage of each pial arteriole in APP23 Tg mice vs. age. Each mouse is depicted with a different symbol (n = 4 Tg mice, 4 pial arterioles per mouse). Line represents LME model with age as the predictor variable, and mouse and vessel as random effects (shaded areas represent the 95% confidence interval of the model’s prediction, Model 1, see Supplemental Table 1). C) Normalized arteriolar diameter vs. age (each arteriolar segment diameter normalized to initial imaging session). Each mouse is depicted with a different symbol (n = 4 Tg, 4 WT mice, 355 measurements across mice). Line represents LME model with best fit including mouse and vessel as random effects, and age, genotype, and age*genotype interactions as fixed effects (shaded areas represent the 95% confidence interval of the model’s prediction, Model 3, see Supplemental Table 2). No change in vessel diameter with age in WT mice, increases in vessel diameter with age in Tg mice, with significant interaction between age*genotype (p = 0.0051). D) Representative examples of arteriole and venule (for reference) RBC velocity calculations from line scan data. E) Normalized arteriolar RBC velocity plotted vs. age (each arteriole segment normalized to first imaging session). Line represents LME model including mouse and vessel as random effects, and age as a fixed effect (shaded areas represent the 95% confidence interval of the model’s prediction, Model 1, see Supplemental Table 3). No significant association with age was observed in this model.

### Pial arteriolar diameter increases with age in APP23 Tg mice, while RBC velocity remains unchanged

Pial arteriolar diameter was observed to increase significantly with age in Tg mice, while remaining unchanged in WT littermates (Figure 1C). LMEs were applied with mouse and vessel as random effects, and age, genotype, and age*genotype interactions as fixed effects, and normalized diameter as the dependent variable. The model with the best fit included all three fixed effects, demonstrating an increase in normalized arteriolar diameter with age*genotype interaction (p = 5.1E-3) (n = 355 vessel measurements across 4 Tg and 4 WT mice). This model explained 26% of the variance in arteriolar diameter (comparison between null and full model: c^2^ = 17.868, p = 4.7E-4). See Supplemental Table 2 for additional model details.

Arteriolar RBC velocity was found to be unchanged with age in both Tg and WT mice (Figure 1D). LMEs were built as described for arteriolar diameter, and no significant relationships between age and/or genotype and RBC velocity were observed (n = 334 measurements across 4 Tg and 4 WT mice). See Supplemental Table 3 for additional model details.

Note, there were no significant differences in baseline arteriolar diameter or baseline arteriolar RBC velocities between WT and Tg mice (students t-test, Supplemental Figure 1).

### Spontaneous vasomotion declines with age in APP23 Tg mice

Spontaneous vasomotion in pial arterioles was observed to decline with age in Tg mice, while remaining unchanged in WT littermates (Figure 2). LMEs were applied with arteriolar vasomotion peak as the dependent variable, mouse and vessel as random effects, and % CAA coverage, age, genotype, and age*genotype interactions as fixed effects. The model with the best fit included all fixed effects, with significant contributions of genotype (p = 2.2E-4) and age*genotype interactions (p = 7.3E-5) (n = 135 measurements over 4 Tg and 4 WT mice). Notably, in this model, % CAA coverage in the imaged FOVs did not have a significant effect (but did have a significant effect in models not including an age*genotype interaction), suggesting that age*genotype interactions may be better predictors of vasomotion decline than local % CAA coverage. This model explained 34% of the variance in the data and fit significantly better than all models tested (all model details in Table 1, comparison between null and full model: c^2^ = 43.126, p = 9.8E-9). No difference was observed in the peak vasomotion frequency between Tg and WT mice, and there was no shift in the peak frequency over the imaging period (Supplemental Figure 3).

**Figure 2.**
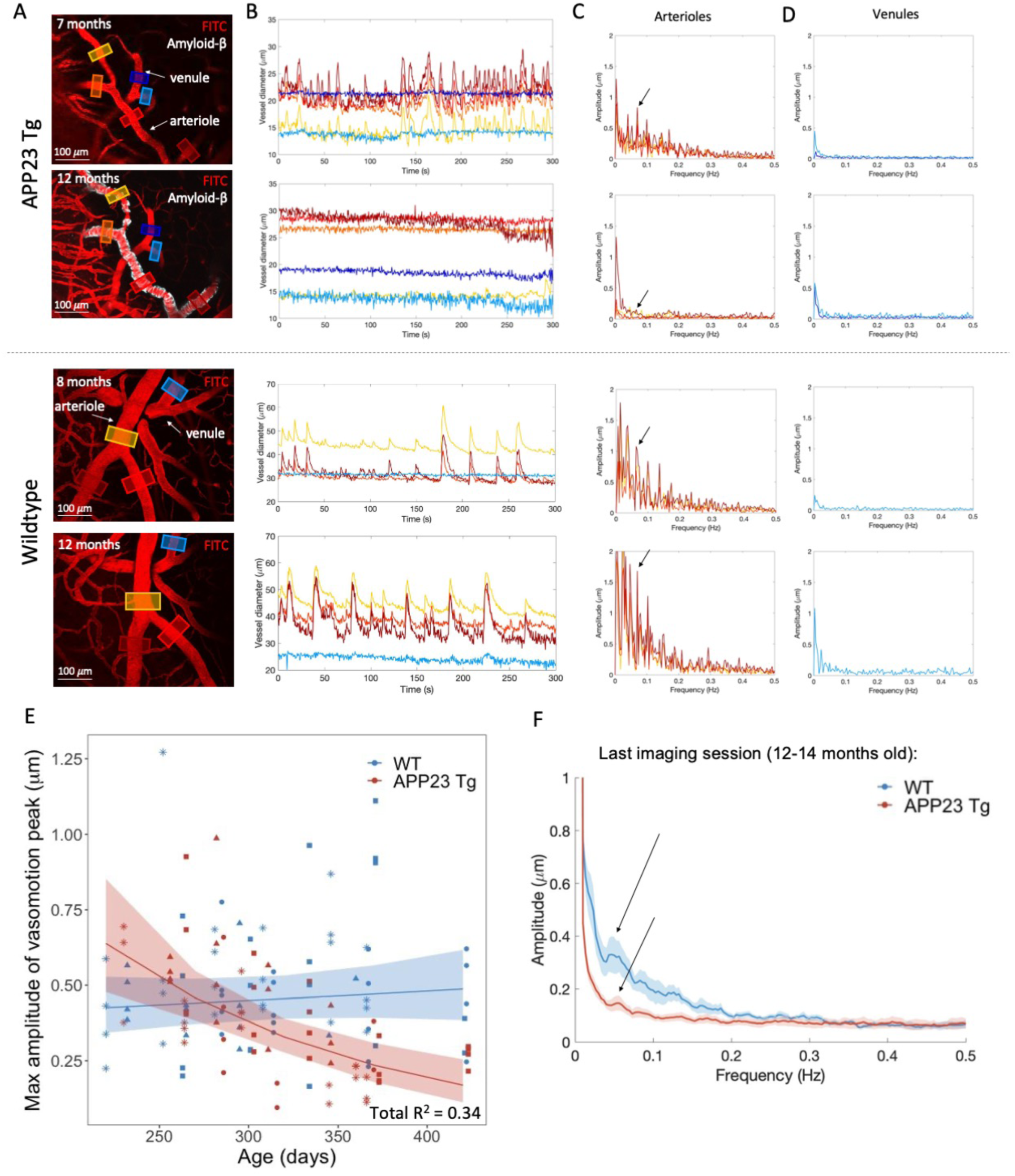
Spontaneous vasomotion decreases significantly with age in APP23 Tg mice. A) Representative examples of APP23 Tg and WT mice at 7 and 12, and 8 and 12 months old respectively. B) Vessel diameter time courses from ROIs depicted in A. Arterioles are observed to have spontaneous fluctuations in vessel diameter, while venules do not notably change in diameter. These fluctuations in vessel diameter are no longer present in Tg mice at 12 months of age, while remaining intact in WT mice at 12 months. C and D) Fast-Fourier transform of time courses shown in B for arterioles (C) and venules (D). Arrows point to the peak detected amplitude between 0.04 and 0.13 Hz, corresponding the “vasomotion peak” frequency. E) Graph of the max amplitude of the vasomotion peak in arterioles vs. age (n = 4 Tg, 4 WT mice, 135 total measurements). Each mouse is represented by a different symbol. LME with the best fit shown, including mouse and vessel as random effects, and % CAA coverage, age, genotype, and age*genotype interactions as fixed effects (shaded areas represent the 95% confidence interval of the model’s prediction, Model 4, see Table 1). Both genotype and age*genotype interactions were significant predictors (p = 2.2E-4 and p = 7.3E-5 respectively). F) Averaged fast-Fourier transforms of time courses from the last imaging sessions from each mouse (n = 4 WT, 4 Tg mice). Mean time courses +/- SEM (shaded) shown. Arrows point to vasomotion peaks in WT and APP23 Tg mice.

To confirm that these changes in vasomotion were not secondary to differential effects of longitudinal cranial windows on Tg mice as compared to WT mice, a separate cohort of mice ∼ 15-16 months of age were imaged awake 1 month following cranial window surgery (n = 4 WT, 3 Tg). We again observed that the maximum amplitude of the vasomotion peak (averaged across each mouse) was significantly lower in the Tg mice than WT littermates (one-way ANOVA with p = 0.0018; post-hoc comparisons using Šídák correction demonstrated significant differences between WT and Tg mice at 12-14 months old (p < 0.01) and 15-16 months old (p < 0.05) with no significant differences at 7-9 months old (Supplemental Figure 2). Notably, the amplitude of the vasomotion peak in the WT mice at 15-16 months of age was similar to the vasomotion peak in the WT mice at 12 months of age, suggesting that vasomotion remains intact at 15-16 months of age in WT mice.

### Beat-to-beat pulsatility increases with age in both APP23 Tg mice and WT littermates

To assess beat-to-beat pulsatility, arteriolar diameter was measured at high temporal resolution using bidirectional line scans (Figure 3). Pulsatility was calculated as both the absolute value of the area under the curve (AUC) of the normalized diameter tracing after detrending (Figure 3C) as well as the peak amplitude of the diameter tracing in the frequency domain (Figure 3D, corresponding to heart rate). Both metrics were observed to increase significantly with age in Tg mice and WT littermates. For AUC comparisons, an LME was built with mouse and vessel as random effects, and diameter, age, genotype, and age*genotype interactions as fixed effects (n = 355 measurements across 4 Tg and 4 WT mice). The model with the best fit (see Supplemental Table 4) demonstrated increases in pulsatility with vessel diameter (p = 3.8E-9), age (p = 1.2E-3), Tg genotype (p = 0.040), and age*genotype interactions (p = 1.2E-3), indicating that pulsatility increases in both Tg and WT mice with age, and that this effect is more prominent in Tg mice. This model explained 48% of the variance in the pulsatility (comparison between null and full model: c^2^ = 102.68, p = 2.2E-16).

**Figure 3.**
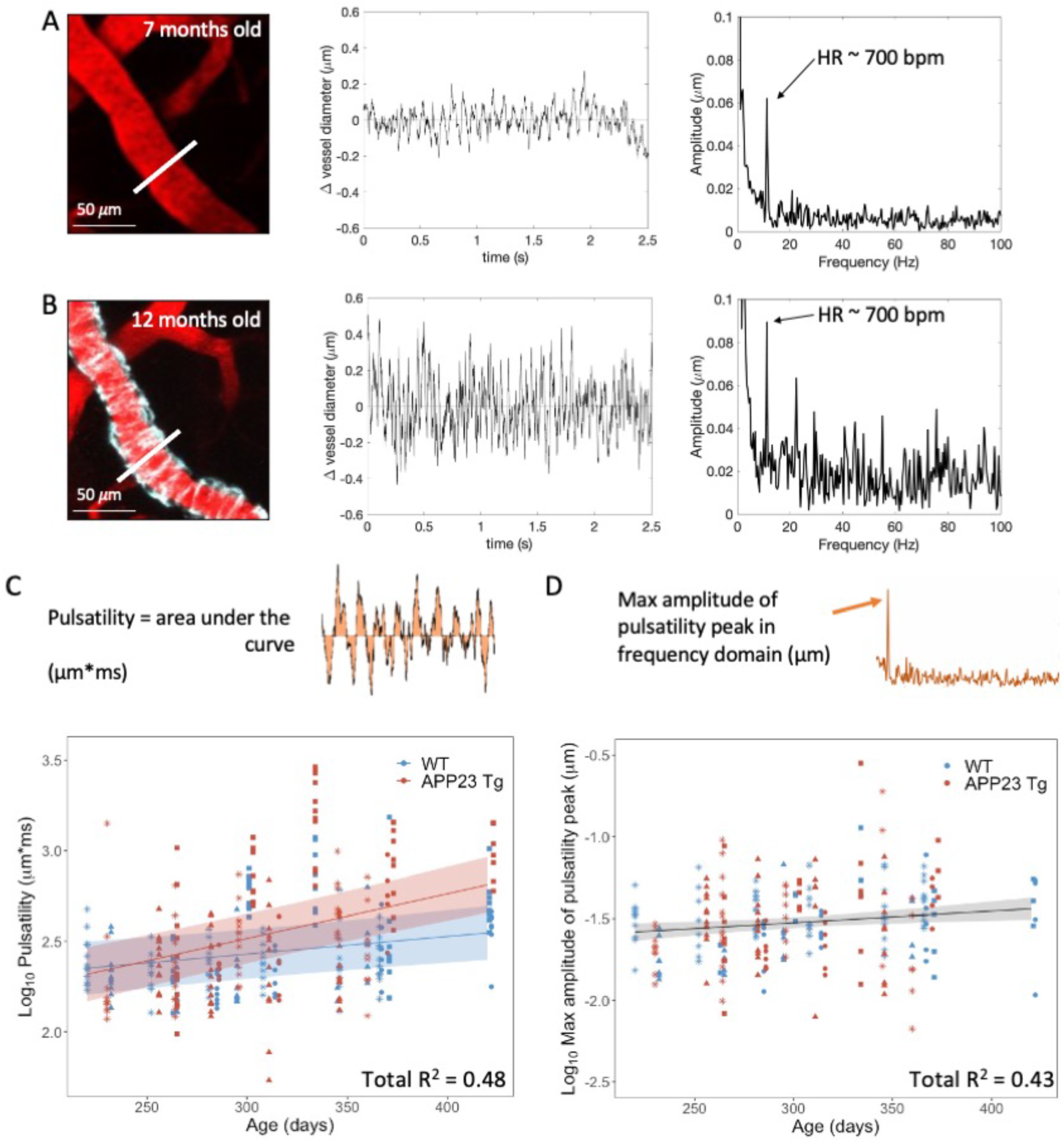
Beat-to-beat pulsatility increases with age in both APP23 Tg and WT mice. A) Representative arteriole in a 7-month-old APP23 Tg mouse captured with *in vivo* 2-photon microscopy (left), vessel diameter time course from rapid line scan (center), and fast-Fourier transform of this time course (right). Arrow pointing to peak frequency, representing heart rate (HR) in beats per minute (bpm). Same measurements from same arteriolar segment approximately 5 months later. C) Log_10_ of pulsatility measurements (calculated as area under the curve, diagram shown) vs. age (n = 4 Tg, 4 WT mice, 355 measurements total). LME with best fit shown, including mouse and vessel as random effects, and diameter, age, genotype, and age*genotype interactions as fixed effects. Increases in pulsatility associated with increasing vessel diameter (p = 3.8E-9), age (p = 0.0012), Tg genotype (p = 0.040), and age*genotype (Tg) interactions (p = 0.0012) (shaded areas represent the 95% confidence interval of the model’s prediction, Model 4, see Supplemental Table 4). D) Log_10_ of pulsatility measurements (calculated as the peak amplitude in the frequency domain, diagram shown) vs. age (n = 4 Tg, 4 WT mice, 242 measurements total). LME with best fit shown, including mouse and vessel as random effects, and diameter and age as fixed effects., demonstrating increases in pulsatility with vessel diameter (p = 4.0E-13) and age (p = 7.7E-3) (shaded areas represent the 95% confidence interval of the model’s prediction, Model 2, see Supplemental Table 5).

To minimize potential effects of noise in the measurements, these data were also analyzed in the frequency domain with a fast-Fourier transform, assessing the amplitude peak at the heart rate frequency. Measurements were excluded (n = 113 measurements across 8 mice) from this analysis if the calculated heart rate peak was > 50% different from the heart rate of the animal (as determined by RBC velocity tracings) as this was suggestive of noise in the data. Again, LMEs were built with the same parameters as above (n = 242 measurements across 4 Tg and 4 WT mice). The model with the best fit (see Supplemental Table 5) included diameter and age (and not genotype or age*genotype interaction), demonstrating increases in pulsatility with vessel diameter (p = 4.0E-13) and age (p = 7.7E-3). This model explained 43% of the variance in the pulsatility (comparison between null and full model: c^2^ = 63.504, p = 1.6E-14). These findings confirm that pulsatility increases with age in Tg and WT mice but suggest that this effect may not be enhanced by Tg genotype.

Note, no significant differences in baseline heart rate were observed between Tg and WT mice, as measured from velocity line scans (Supplemental Figure 1). Additionally, no significant changes in heart rate were observed with increasing age in either Tg or WT mice (Supplemental Figure 4, Supplemental Table 6).

### Baseline CBF is decreased in APP23 Tg mice at 24 months of age

Separate cohorts of mice were imaged with *in vivo* 9.4T MRI to capture global CBF and CVR in Tg and WT mice at advanced ages. Figure 4 displays averaged CBF and CVR maps across the age groups (12, 18, and 24 months) for Tg and WT mice in a single brain slice. The results for the other four brain slices are shown in Supplemental Figure 5. Globally decreased baseline CBF was observed in the maps of the 18- and 24-month-old (but not the 12-month-old) Tg as compared to WT mice. Figure 5a shows the absolute and baseline-corrected group-averaged CBF time-profiles (mean ± SD) retrieved in the somatosensory cortex. Baseline CBF was significantly lower in 24-month-old Tg (86 mL/100 g/min) as compared to WT mice (109 mL/100 g/min, Z = 2.77, p = 0.0056). No significant reduction was found in the 12- or 18-month-old cohorts, but there was a trend towards a lower median CBF at 18 months (72 mL/100 g/min in Tg vs 106 mL/100 g/min in WT mice, Z = 1.91, p = 0.056).

**Figure 4.**
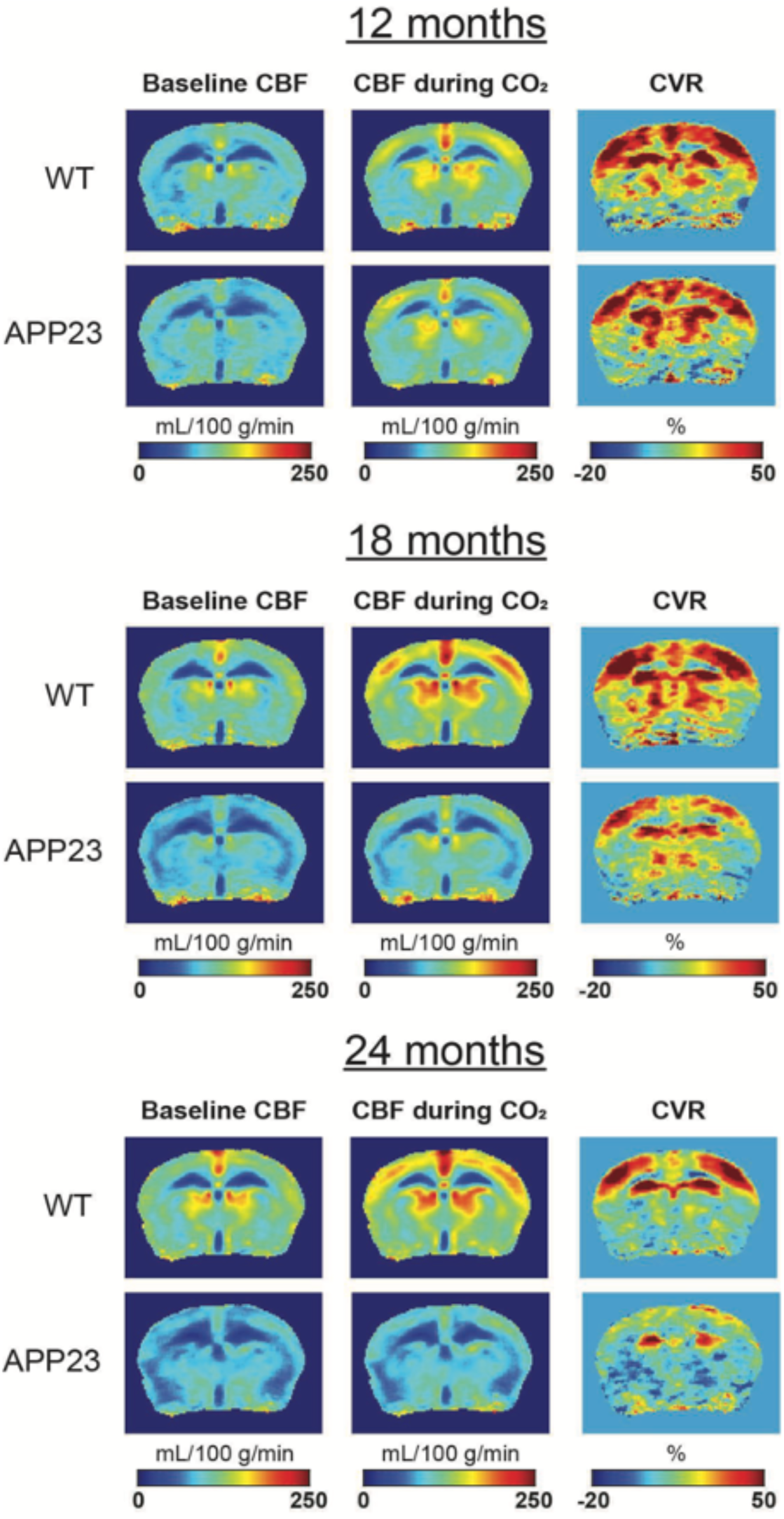
MRI maps of cerebral blood flow (CBF) and cerebrovascular reactivity (CVR). The rows display maps averaged per genotype (wild type [WT] or APP23 transgenic [Tg]) and per age group (12, 18 or 24 months old), with each group consisting of n = 5-9 mice. On the columns, the maps display the baseline CBF, CBF during CO_2_, and CVR (relative CBF change during CO_2_). Note the similarity of the WT and APP23 Tg maps at 12 months of age, and the visibly lower CBF and CVR at 24 months of age. Also note that only the middle brain slice is shown, the results of the other 4 slices are shown in Supplemental Figure 5.

**Figure 5.**
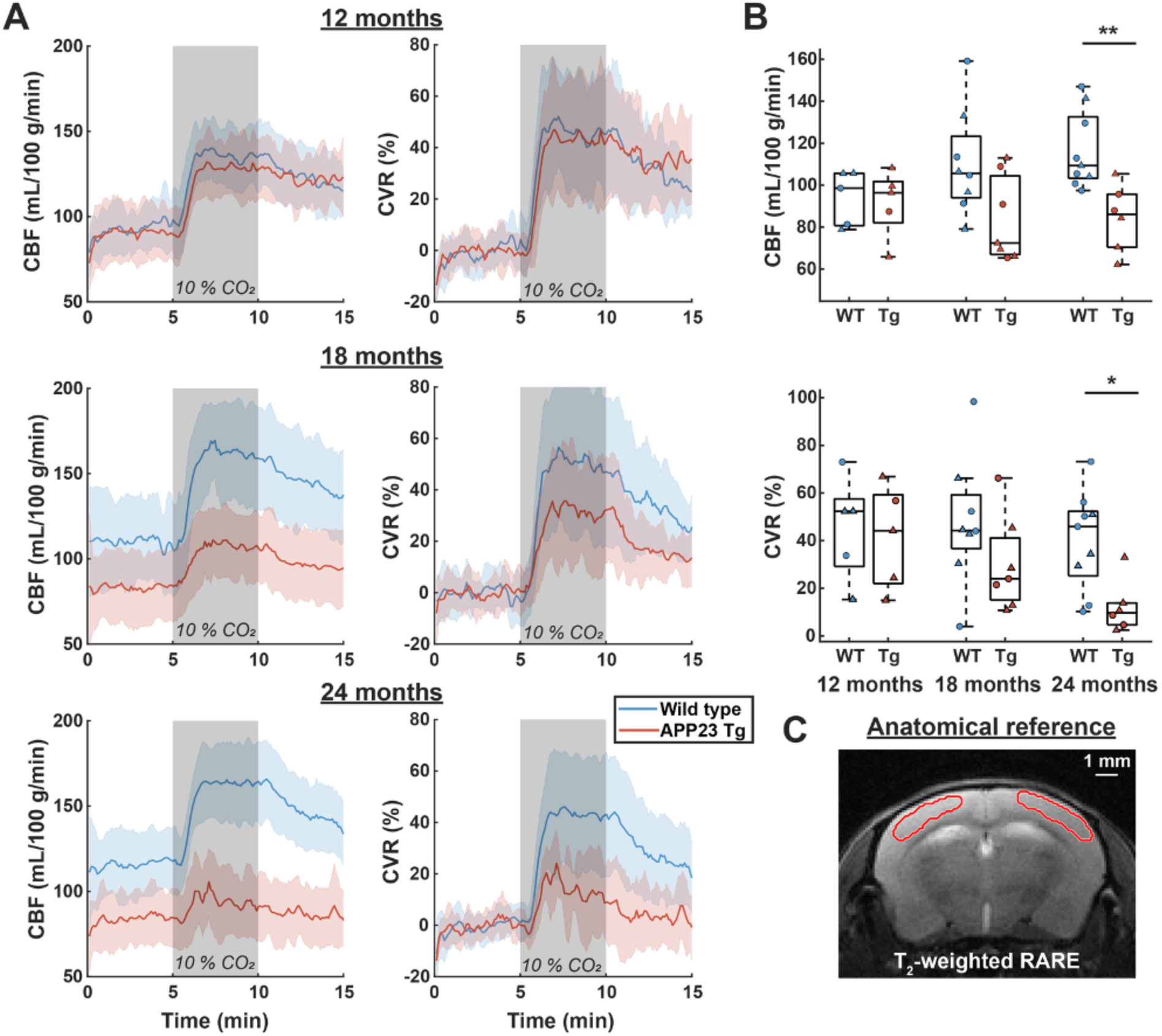
Cerebral blood flow (CBF) and cerebrovascular reactivity (CVR) values in the somatosensory cortex. In the left column of A), the average CBF time-profiles (± standard deviation [SD]) are shown, per genotype (wild type [WT] in blue, APP23 transgenic [Tg] in red) and per age group, with each group consisting of n = 5-9 mice. The right column displays CVR time-profiles, i.e., baseline-corrected CBF time-profiles. In B), boxplots are shown per genotype and age-group, where every point represents the average CBF (top) or CVR (bottom) per mouse. Dots represent male and triangles represent female mice. CBF was significantly lower in the 24-month-old APP23 Tg mice when compared to WT mice (Z = 2.77, p = 0.0056, Mann Whitney U test). CVR was also significantly lower in the 24-month-old APP23 Tg mice (Z = 2.42, p = 0.016, Mann Whitney U test). In C), the red outline displays the somatosensory cortical region that was analyzed, overlaid on the reference T2-weighted RARE scan. * p < 0.05, ** p < 0.01.

### CVR in response to a CO_2_ challenge is decreased in APP23 Tg mice by 24 months of age

The CVR maps demonstrate differences in CVR between WT and Tg groups predominantly in the somatosensory cortex in the 18- and 24-month-old cohorts (but not the 12-month-old) (Figure 4). In the 24-month-old Tg mice, CVR was significantly lower as compared to WT mice (10% in Tg vs 47% in WT, Z = 2.42, p = 0.016), and non-significant trends for lower CVR in the 18-month-old Tg mice were also observed (25% in Tg vs 45% in WT, Z = 1.22, p = 0.22) (Figure 5).

Visual inspection of angiograms acquired with TOF did not demonstrate abnormalities in the larger arteries of Tg mice (Supplemental Figure 6). Furthermore, no differences were found in the labeling efficiency between the Tg and WT mice (Supplemental Figure 7), nor in the magnitude of change in transcutaneous pCO_2_ following the CO_2_ challenge (Supplemental Figure 8).

### Vasodilatory response to isoflurane is reduced in APP23 Tg mice

Given that the 2-photon experiments demonstrated an increase in baseline pial arteriolar diameter in Tg mice with age (and no change in RBC velocity), one might have expected an increase rather than a decrease in CBF in the Tg mice as compared to WT mice. One potential explanation for the apparent discrepancy between these results is a differential response to the light isoflurane anesthesia (1.1%) used in the MRI experiments. Note, all 2-photon imaging was performed awake. We hypothesized that arterioles in Tg mice would not dilate significantly in response to isoflurane, potentially leading to relatively lower pCASL-based baseline CBF as compared to WT mice. To test this hypothesis, we performed a separate set of 2-photon experiments in 15-month-old Tg and WT mice (n = 3 WT, 3 Tg) in which we systematically increased the level of anesthesia from awake to 0.5%, 1.1%, 1.5%, and 2% isoflurane. As expected, we observed large increases in vessel diameter in 2/3 WT mice. In comparison, we observed smaller increases in vessel diameter in 2/3 of the Tg mice (Supplemental Figure 9). Given these findings, it remains unclear whether baseline CBF is also decreased in Tg mice under awake conditions.

### APP23 Tg mice develop cortical microbleeds between 12 and 18 months of age

The same mice used for CBF and CVR measurements were used to assess MRI-visible microbleeds. Cortical microbleeds were rated on MGE sequences (Figure 6A). At 12 months of age, microbleeds were observed in 2/5 of the Tg mice (range 0-2). At 12 months of age, microbleeds were observed in 5/7 of the Tg mice (range 0-2). The highest number of microbleeds was observed at 24 months of age (in 5/6 mice, range 0-7). Across all three cohorts, the number of microbleeds inversely correlated with CVR (Spearman’s ρ = -0.506, p = 0.032), but not with CBF (Spearman’s ρ = -0.044, p = 0.863). Note, only cortical microbleeds were assessed, as hypointensities on MGE sequences in deeper structures of the brain are less specific and may represent mineralization, which has been previously observed in aged WT mice^26^.

**Figure 6.**
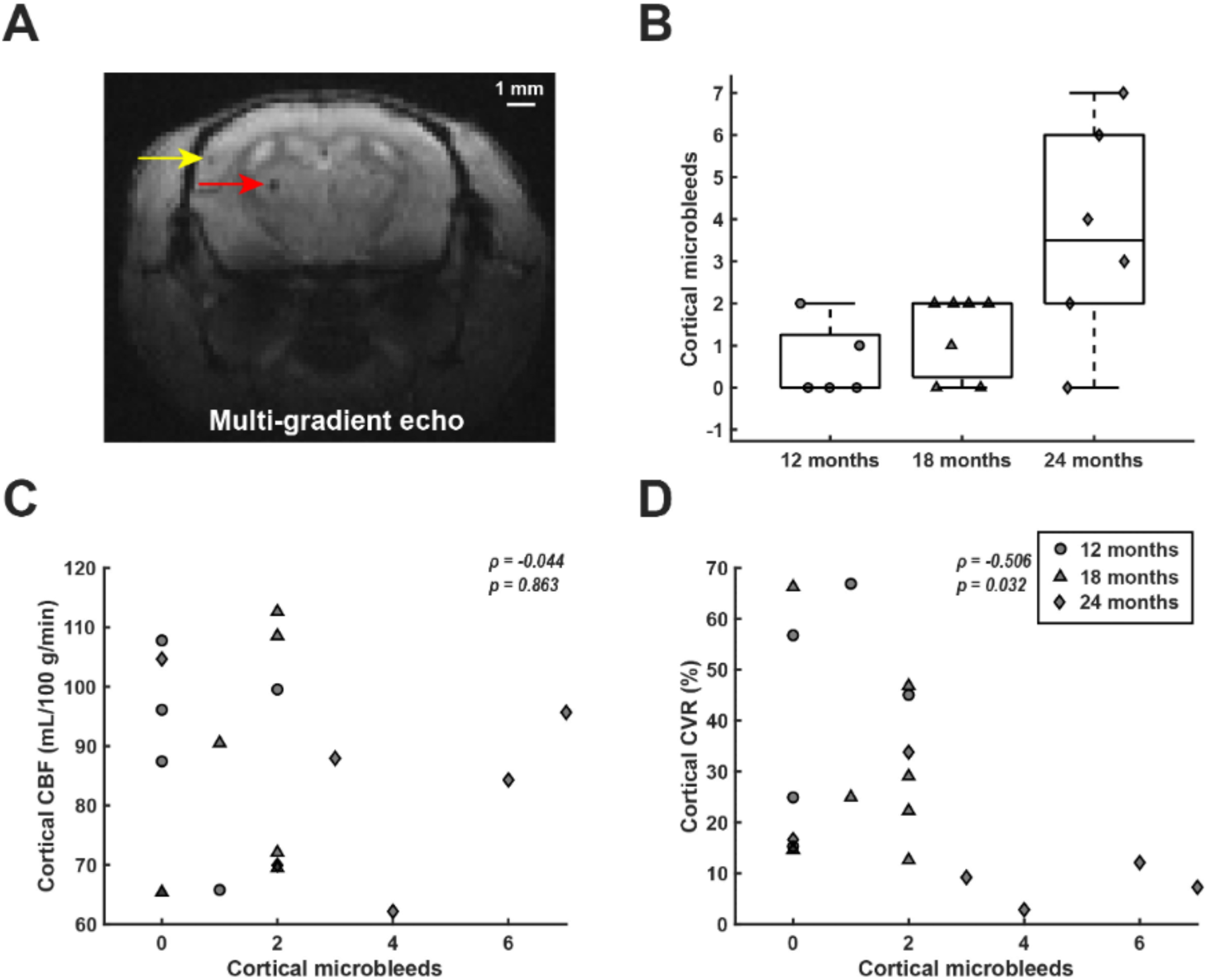
Microbleed quantification and correlation with cerebral blood flow (CBF) and cerebrovascular reactivity (CVR) in APP23 Transgenic (Tg) mice. Microbleeds were scored in APP23 Tg mice by counting hypointense round or ovoid lesions in the cortex on multi-gradient echo (MGE) scans, as illustrated by the yellow arrow in A). Note that sub-cortical hypointensities (red arrow) were not scored. B) shows the number of cortical microbleeds per age-group. In C), the number of microbleeds over all age-groups is plotted against cortical CBF, which did not reveal a correlation (Spearman’s ρ = -0.044, p = 0.863). Circles, triangles and diamonds represent 12, 18, and 24-month-old mice respectively. In D), the number of cortical microbleeds over all age-groups is plotted against cortical CVR, which revealed a significant correlation (Spearman’s ρ = -0.506, p = 0.032).

### Arteriolar smooth muscle actin is decreased in 18- and 24-month-old APP23 Tg mice

A subset of mice was selected for immunohistochemistry including staining of serial sections for amyloid-β and SMA (see Methods). Pial arterioles were rated on both the Vonsattel grade of individual vessel CAA severity and SMA score (blinded to age and other stains, Figure 7A). Note, few arterioles were observed to be grade 0 (no amyloid-β) in all Tg mice, and low numbers of Grade 1 (incomplete amyloid-β coverage) were observed in 18-month and 24-month-old Tg mice (Figure 7B). Most vessels in 18-month and 24-month-old age groups were grade 2 (circumferential amyloid-β). No amyloid-β was observed in WT animals (all “grade 0” vessels). SMA scores for all arterioles of each grade assessed were averaged within each mouse to generate vessel grade specific SMA scores (Figure 7C).

**Figure 7.**
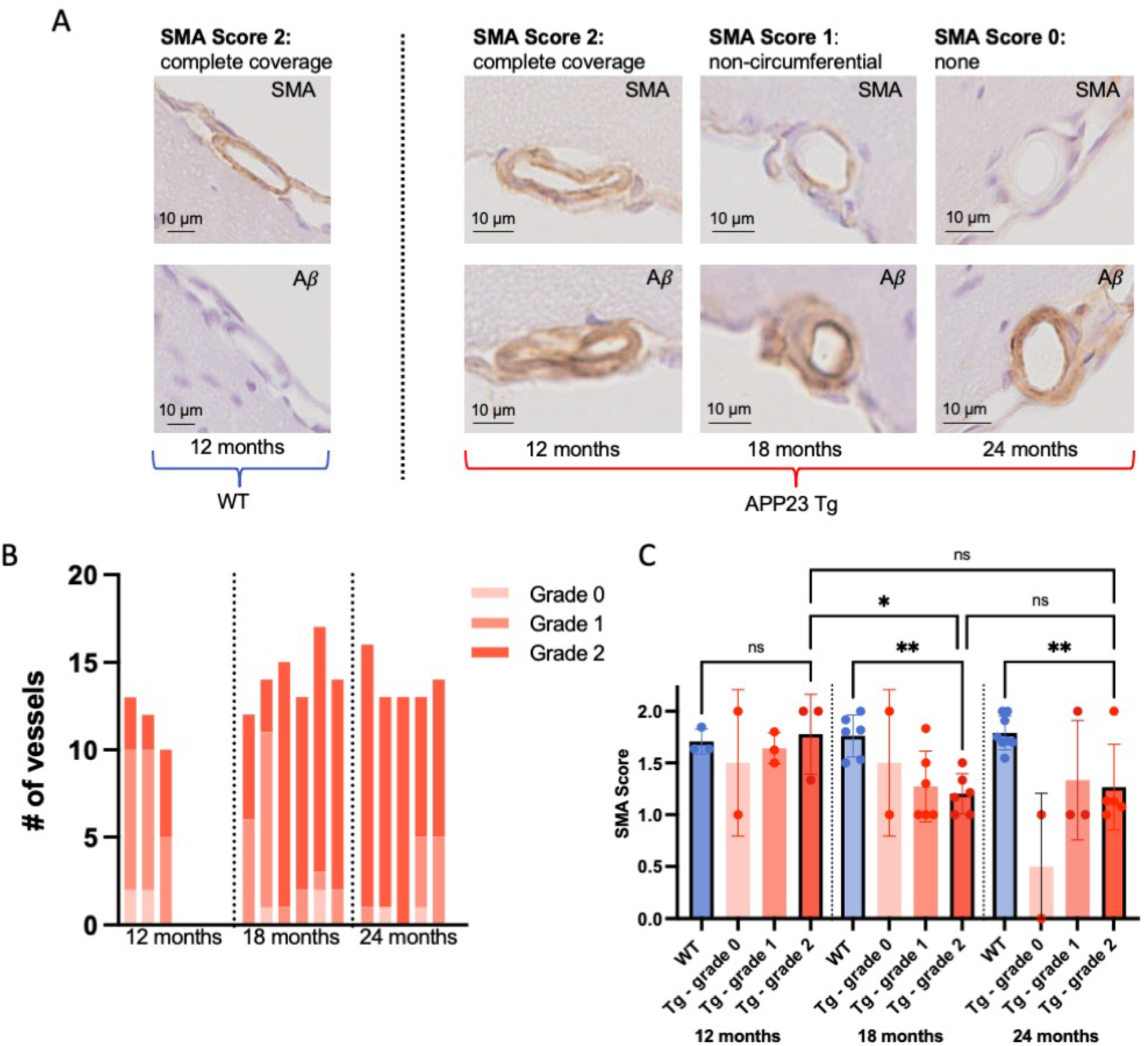
Smooth muscle actin is decreased in arterioles with vascular amyloid-β in 18- and 24-month-old APP23 Tg mice. A) SMA and amyloid-β stains from a representative SMA score 2 vessel from a WT mouse and representative SMA score 2, 1, and 0 vessels from APP23 Tg mice. B) Numbers of Vonsattel grade 0, 1, and 2 pial arterioles identified in APP23 Tg mice of different ages. Each column represents a separate mouse (1 brain section analyzed per mouse). C) Average pial vessel SMA score per mouse. For APP23 Tg mice, only SMA scores from Vonsattel grade 2 vessels were used for statistical analysis. One-way ANOVA demonstrated significant differences across these groups (p = 0.0005), pre-selected comparisons performed using Šídák correction shown (* p < 0.05, ** p < 0.01).

Given the low numbers of grade 0 and grade 1 vessels in Tg mice across the age groups, only grade 2 vessels were included in formal statistical analysis of SMA coverage. SMA scores from all vessel grades are shown in Figure 7C. An ordinary one-way ANOVA demonstrated significant differences across these groups (p = 0.0005), and a post-hoc analysis was performed on pre-selected comparisons using a Šídák correction. SMA coverage in grade 2 pial arterioles was observed to be significantly lower in 18-month-old Tg mice as compared to 12-month-old Tg mice (p < 0.05, Figure 7). Additionally, while there was no significant difference between SMA coverage in 12-month-old WT arterioles vs. 12-month-old Tg grade 2 arterioles, significant differences were observed between 18-month-old WT and Tg (p < 0.01), and 24-month-old WT and Tg mice (p < 0.01).

## DISCUSSION

In this work, we assessed several measures of vascular structure and function in the APP23 mouse model of amyloidosis, using both *in vivo* 2-photon microscopy and MRI. An early loss of spontaneous vasomotion at the level of individual arterioles was detected with 2-photon microscopy, occurring on a similar timeline as the appearance of vascular amyloid-β (at ∼8 – 10 months of age). Baseline arteriolar diameter also increased during this age range. Given VSMC are essential to both vasomotion^8,9^ and maintaining cerebrovascular tone^27^, these results suggest VSMC dysfunction begins to occur at the time of initial amyloid-β deposition.

Prior studies have suggested that spontaneous vasomotion and functional hyperemia, both dependent on VSMC function, are important for glymphatic clearance (including amyloid-β) from the brain^9,10^. Given initial amyloid-β deposition is thought to occur approximately 30 years prior to the development of symptoms in patients with CAA^28^, the identification of vasomotion as an early biomarker for VSMC dysfunction, preceding VSMC loss, may provide a critical treatment window for interventions targeted at enhancing vasomotion or neurovascular coupling to drive glymphatic clearance. We observed that despite evidence of VSMC dysfunction, CVR (as measured by the response to inhaled CO_2_) was not yet altered at 12 months of age in Tg mice. Additionally, we observed that VSMCs expressing SMA (a contractile protein) were still present in pial arterioles with circumferential amyloid-β in 12-month-old Tg mice. Similarly, a prior study of APP/PS1 mice demonstrated that while global amyloid-β burden was associated with VSMC loss, local amyloid-β deposition was not tightly linked to VSMC loss at an individual vessel level^9^, suggesting a potential role for other factors such as soluble amyloid-β in VSMC dysfunction/loss. Further supporting this observation, in the vasomotion LME model with the best fit, we found that the % local CAA coverage did not have a significant effect on vasomotion (but did have an effect in models not including an age*genotype interaction). This finding suggests again that there may be other, potentially more global, factors driving the observed local decline in vasomotion. However, we note that a prior study of Tg2576 mice (another model of amyloidosis) at 8 months of age did not demonstrate any impairments in VSMC function despite the presence of increased soluble amyloid-β, and VSMC function was subsequently impaired in older mice with vascular amyloid-β deposition^29^.

Importantly, our findings suggest that some components of the microvasculature maintain the ability to vasodilate despite initial amyloid-β deposition. Therefore, interventions aiming to enhance/drive vascular reactivity may still be effective despite the loss of spontaneous vasomotion. However, a limitation to this study is that we were unable to determine which individual vessels were dilating in response to a CO_2_ challenge in the MRI portion of the study. Therefore, it remains a possibility that these findings reflect intact vasodilation at the capillary/diving arteriolar level, while the loss of vasomotion we observed with 2-photon microscopy was at the pial arteriolar level. Of note, a prior study from our group of 8–10-month-old APP/PS1 mice (another model of amyloidosis) demonstrated intact vasomotion with impaired vascular reactivity in response to visual stimuli^9^. Future studies are needed to determine whether and when sensory stimulus-evoked vascular reactivity is affected in APP23 mice; it is possible that pathways dependent on neurovascular coupling are more profoundly affected by vascular (and parenchymal) amyloid-β accumulation.

We also observed increases in beat-to-beat pulsatility of the arterioles with age in both WT and Tg mice, increases which were potentially more pronounced in Tg mice. A more pronounced increase in pulsatility with age has also been previously observed in APP/PS1 mice^30^. Increases in beat-to-beat pulsatility in small vessels with age have also been observed in human subjects^31^, irrespective of the presence of small vessel disease^15^. These increases in pulsatility may reflect stiffening of the more upstream vessels (e.g. aorta) with aging^32^, possible increased compliance in the cerebral vasculature, another marker of VSMC dysfunction, and/or decreases in brain tissue stiffness with age^33^. While prior findings have suggested a link between increased arteriolar pulsations and glymphatic flow^14^, a study in a mouse model of hypertension demonstrated that alterations/increases in vessel wall pulsatility may in fact disrupt perivascular clearance^34^.

Reduced vasomotion occurred before the appearance of microbleeds on MRI (between 12 and 18 months) and before CBF and CVR reductions were detectable with MRI (between 18 and 24 months). Notably, a reduction of CBF has been previously reported in APP23 mice at a similar age^35^, and the occurrence of microbleeds between 12 and 18 months of age is also in line with data of others^17,36^. The reduced CVR in response to the strong CO_2_ stimulus in 24-month-old APP23 Tg mice is particularly noteworthy, given the lower baseline CBF in this group. As CVR is measured relative to baseline CBF, one would expect confounds due to a lower baseline CBF to result in an enhanced relative CVR, rather than a diminished CVR as was observed. A significant correlation was found between microbleed count and CVR; however, it is unclear whether these are causally related or just independent results of the CAA pathology.

The findings of increases in baseline arteriolar diameter with age as measured with 2-photon microscopy together with a CBF decrease as measured with MRI are counter-intuitive. Vessel diameter increases with no change in RBC velocity would typically be expected to lead to increased CBF. Notably, local vasodilation caused by fibrillar amyloid-β on pial arteries, as observed here, has been previously reported^37^. However, some studies have shown that amyloid-β causes vasoconstriction^38,39^, including a recent study of dissected and pressurized pial arteries from 18-month-old APP23 Tg mice. The different findings could possibly be explained by the different forms of amyloid-β studied, i.e. soluble or fibrillar forms of amyloid-β, or by different vessel types affected, i.e. pial or penetrating arteries or capillaries, or by differences in experimental preparation. It remains possible that *in vivo* in APP23 Tg mice, at sites further down the vascular tree, soluble amyloid-β and/or fibrillar amyloid-β may act as a vasoconstrictor. Given these sites are at a point of higher resistance in the vascular tree, these effects may dominate CBF measurements leading to our observed reduction in CBF. Alternative or additive explanations for lower observed CBF in APP23 mice include decreased vasodilation of arterioles in APP23 Tg mice in response to isoflurane anesthesia, which was used for the MRI experiments but not for the 2-photon experiments. Similar findings of decreased vasodilation of isoflurane anesthesia have been observed in Tg2576 mice at advanced ages (19 months)^29^. This differential response to isoflurane led to relatively increased CBF measurements in WT mice as observed via laser doppler in this study of Tg2576 mice^29^. Other considerations may be reduced capillary density and/or reduced metabolism in APP23 Tg mice. Indeed, reduced capillary density has been reported in APP23 mice^40^, and the high burden of parenchymal amyloid-β plaques observed in the model at advanced ages may impact neuronal activity. In general, baseline CBF is not considered a sensitive marker of CAA, as CBF has been observed to be unchanged in some studies^6^, or only mildly reduced in others^7,41^. Interestingly however, in the healthy elderly population, the presence of cortical cerebral microbleeds has been associated with lower CBF^42^.

An unusual finding in this study is the increase in baseline CBF in WT mice with age. This may be an artifact related to a batch effect, since all the 12-month-old animals were bred and aged at the Jackson laboratories, and the other age groups were a mix of home-bred and Jackson lab bred. It is possible that stress during transportation led to endothelial dysfunction in these mice, which has been previously reported^43,44^. This could then lead to reduced vasodilation upon isoflurane administration, and thus lower CBF in the Jackson mice, i.e. in the younger mice of our MRI cohort. Importantly however, between genotypes, the ratio of mice derived from the Jackson lab or in-house was the same, indicating that the CBF and CVR reductions in Tg mice are not biased by batch effects.

There are several limitations in this study. The group sizes in the different cohorts were relatively small and sex was not always equally distributed among the groups. However, no obvious separation was apparent between females and males in the assessed measures of vascular function, although prior studies have noted a higher amyloid-β plaque load (both vascular and parenchymal) in female APP23 mice^45^. Of note, the Tg mouse in which we observed no amyloid-β plaques throughout the study was male. Another limitation is the cranial window surgery in the 2-photon cohort, as this may have impacted vascular function. These mice were allowed to recover for at least 4 weeks post-surgery prior to their initial imaging sessions, with a goal of minimizing the acute effects of the surgery on neurovascular function. Additionally, we did not control for potential confounders such as behavior, state of arousal, and locomotion which have been shown to affect neurovascular coupling and may also impact vasomotion^46–48^. Lastly, the use of isoflurane in the MRI cohort may have influenced the results. However, the fact that a combination of imaging modalities was used in this study, each with a different set of limitations, strengthens the findings, and underscores that vascular dysfunction is a prominent early feature of the APP23 mouse model.

To conclude, this study found an early loss of vasomotion in APP23 Tg mice, occurring at approximately the same time as vascular amyloid-β deposition, before the loss of VSMCs, reduction of CVR, and occurrence of cortical microbleeds. Detection of these changes in vasomotion may present an opportunity for early, pre-symptomatic diagnosis of CAA. Additionally, given the importance of vasomotion for glymphatic amyloid-β clearance, preserving vasomotion in the pre-symptomatic phase of the disease may be an important target for prevention studies in CAA. Future studies should be considered to determine whether this early loss of vasomotion may potentially precede initial amyloid-β deposition.

## Abbreviations

CAA: cerebral amyloid angiopathy
ICH: intracerebral hemorrhage
VSMC: vascular smooth muscle cells
CSF: cerebrospinal fluid
MGE: multi-gradient echo
pCASL: pseudo-Continuous Arterial Spin Labeling
Tg: transgenic
WT: wildtype
CBF: cerebral blood flow
CVR: cerebrovascular reactivity
TOF: time-of-flight
SMA: smooth muscle actin
MRI: magnetic resonance imaging
BOLD: blood oxygen level dependent
TR: repetition time
TE: echo time
RARE: Rapid Acquisition with Relaxation Enhancement
EPI: echo-planar imaging
fc-FLASH: flow-compensated, fast low angle shot
RBC: red blood cell
FOV: field of view
ROI: region of interest
PBS: phosphate buffered saline
TBS: tris buffered saline
DAB: 3,3’-diaminobenzidine
LME: linear mixed effects
AIC: Akaike information criterion
BIC: Bayesian information criterion
AUC: area under the curve

## Acknowledgements

The authors thank Drs. Orla Bonnar, Valentina Perosa, and Maria Sanchez Mico for helpful discussions. The authors also thank Drs. Joseph Mandeville and Andre van der Kouwe for their help with setting up the MRI protocol.

## Supplemental figure captions

**Supplemental Figure 1.** N**o difference in baseline arteriolar diameter, velocity, or heart rate between APP23 Tg mice and WT littermates.** Mean +/- SD shown. Left: n = 39, 42 arterioles; middle: n = 43, 42 arterioles; right: n = 43, 42 arterioles (note, each measurement treated as independent).

**Supplemental Figure 2. Vasomotion remains intact in a separate group of WT mice at 15-16 months of age.** The same cohort of mice was imaged between 7-14 months (n = 4 WT, 4 Tg). The 15–16-month-old mice shown represent an independent cohort of mice (n = 4 WT, 3 Tg). One-way ANOVA (p = 0.0018), pre-selected comparisons performed using Šídák correction shown (* p < 0.05, ** p < 0.01).

**Supplemental Figure 3. No difference in vasomotion frequency between APP23 Tg and WT mice.** Frequency does not change over the imaged age range (n = 4 WT, 4 Tg mice).

**Supplemental Figure 4. No change in heart rate with age in APP23 Tg and WT mice.** Each mouse is depicted with a different symbol (n = 4 Tg, 4 WT mice, 358 measurements across mice). Line represents LME model including mouse and vessel as random effects and age as a fixed effect (shaded areas represent the 95% confidence interval of the model’s prediction, Model 1, see Supplemental Table 6). No significant association with age was observed in this model.

**Supplemental Figure 5. Whole-brain MRI maps of cerebral blood flow (CBF) and cerebrovascular reactivity (CVR).** The rows display maps averaged per genotype (wild type [WT] or APP23 transgenic [Tg]) and per age group (12, 18 or 24 months old), with each group consisting of n = 5-9 mice. On the columns, the maps display the baseline CBF, CBF during CO_2_, and CVR (relative CBF change during CO_2_), and in the sub-columns the different 5 MRI slices are shown.

**Supplemental Figure 6. MRI time-of-flight (TOF) images of all mice in the 24-month-old cohort.** Maximum intensity projections of vessel enhancement filtered TOF images are shown for wild type (WT) mice on the top row, and for APP23 transgenic (Tg) mice on the bottom row.

**Supplemental Figure 7. Pseudo-continuous arterial spin labeling (pCASL) inversion efficiency measurements.** Phase and magnitude images of the pCASL fc-FLASH sequence acquired at the level of the carotids are shown in A), both for label and control acquisitions. The relative complex signal difference between label and control images is shown on the bottom image in A), where the right and left carotids are indicated with black arrows. The plot in B) displays the inversion efficiency values measured in the carotids for the 12, 18, and 24-month-old cohorts, showing no significant differences between wild type (WT) and APP23 transgenic (Tg) mice. Dots represent male and triangles represent female mice.

**Supplemental Figure 8. Transcutaneous pCO_2_ (tc-pCO_2_) measurements.** The graphs show the mean mmHg change in tc-pCO_2_ (± standard deviation) during a 10% CO_2_ challenge, acquired in 3 wild type and 3 APP23 transgenic (Tg) mice from the 18-month-old cohort, which underwent a separate 10% CO_2_ challenge outside the MRI scanner.

**Supplemental Figure 9. Baseline arteriolar diameter increases more strongly with isoflurane in 15-month-old WT compared to APP23 Tg mice.** Left: measurements of same arterioles imaged in APP23 Tg and WT mice with increasing isoflurane administered. Right: % change in vessel diameter from awake imaging to 1.1% isoflurane in APP23 Tg and WT mice.

## References

1. van Asch CJ, Luitse MJ, Rinkel GJ, van der Tweel I, Algra A, Klijn CJ. Incidence, case fatality, and functional outcome of intracerebral haemorrhage over time, according to age, sex, and ethnic origin: a systematic review and meta-analysis. Lancet neurology. 2010;9:167–76.

2. Kozberg MG, Perosa V, Gurol ME, van Veluw SJ. A practical approach to the management of cerebral amyloid angiopathy. International Journal of Stroke. 2021;16:356–369.

3. Vonsattel JPG, Myers RH, Tessa Hedley-Whyte E, Ropper AH, Bird ED, Richardson EP. Cerebral amyloid angiopathy without and with cerebral hemorrhages: A comparative histological study. Annals of Neurology. 1991;30:637–649.

4. Love S, Chalmers K, Ince P, Esiri M, Attems J, Jellinger K, Yamada M, McCarron M, Minett T, Matthews F, Greenberg S, Mann D, Kehoe PG. Development, appraisal, validation and implementation of a consensus protocol for the assessment of cerebral amyloid angiopathy in post-mortem brain tissue. American Journal of Neurodegenerative Diseases. 2014;3:19–32.

5. Kozberg MG, Yi I, Freeze WM, Auger CA, Scherlek AA, Greenberg SM, van Veluw SJ. Blood-brain barrier leakage and perivascular inflammation in cerebral amyloid angiopathy. Brain Communications. 2022;4:1–13.

6. Dumas A, Dierksen GA, Gurol ME, Halpin A, Martinez-Ramirez S, Schwab K, Rosand J, Viswanathan A, Salat DH, Polimeni JR, Greenberg SM. Functional MRI Detection of Vascular Reactivity in Cerebral Amyloid Angiopathy. Annals of neurology. 2012;72:997–1003.

7. van Opstal AM, van Rooden S, van Harten T, Ghariq E, Labadie G, Fotiadis P, Gurol ME, Terwindt GM, Wermer M, van Buchem MA, Greenberg SM, van der Grond J. Cerebrovascular function in pre-symptomatic and symptomatic individuals with hereditary cerebral amyloid angiopathy: a case-control study. Lancet Neurol. 2017;16:115–122.

8. Hill RA, Tong L, Yuan P, Murikinati S, Gupta S, Grutzendler J. Regional Blood Flow in the Normal and Ischemic Brain Is Controlled by Arteriolar Smooth Muscle Cell Contractility and Not by Capillary Pericytes. Neuron. 2015;87:95–110.

9. van Veluw SJ, Hou SS, Calvo-Rodriguez M, Arbel-Ornath M, Snyder AC, Frosch MP, Greenberg SM, Bacskai BJ. Vasomotion as a Driving Force for Paravascular Clearance in the Awake Mouse Brain. Neuron. 2020;105:549–561.e5.

10. Holstein-Rønsbo S, Gan Y, Giannetto MJ, Rasmussen MK, Sigurdsson B, Beinlich FRM, Rose L, Untiet V, Hablitz LM, Kelley DH, Nedergaard M. Glymphatic influx and clearance are accelerated by neurovascular coupling. Nat Neurosci. 2023;26:1042–1053.

11. Kedarasetti RT, Turner KL, Echagarruga C, Gluckman BJ, Drew PJ, Costanzo F. Functional hyperemia drives fluid exchange in the paravascular space. Fluids and Barriers of the CNS. 2020;17:1–25.

12. Greenberg SM, Bacskai BJ, Hernandez-Guillamon M, Pruzin J, Sperling R, van Veluw SJ. Cerebral amyloid angiopathy and Alzheimer disease — one peptide, two pathways. Nature Reviews Neurology. 2020;16:30–42.

13. Van Etten ES, Verbeek MM, Van Der Grond J, Zielman R, Van Rooden S, Van Zwet EW, Van Opstal AM, Haan J, Greenberg SM, Van Buchem MA, Wermer MJH, Terwindt GM. β-Amyloid in CSF: Biomarker for preclinical cerebral amyloid angiopathy. Neurology. 2017;88:169–176.

14. Iliff JJ, Wang M, Zeppenfeld DM, Venkataraman A, Plog BA, Liao Y, Deane R, Nedergaard M. Cerebral Arterial Pulsation Drives Paravascular CSF–Interstitial Fluid Exchange in the Murine Brain. J Neurosci. 2013;33:18190–18199.

15. Perosa V, Arts T, Assmann A, Mattern H, Speck O, Oltmer J, Heinze H-J, Düzel E, Schreiber S, Zwanenburg JJM. Pulsatility Index in the Basal Ganglia Arteries Increases with Age in Elderly with and without Cerebral Small Vessel Disease. American Journal of Neuroradiology. 2022;43:540– 546.

16. Winkler DT, Bondolfi L, Herzig MC, Jann L, Calhoun ME, Wiederhold KH, Tolnay M, Staufenbiel M, Jucker M. Spontaneous hemorrhagic stroke in a mouse model of cerebral amyloid angiopathy. The Journal of neuroscience : the official journal of the Society for Neuroscience. 2001;21:1619– 1627.

17. Reuter B, Venus A, Heiler P, Schad L, Ebert A, Hennerici MG, Grudzenski S, Fatar M. Development of Cerebral Microbleeds in the APP23-Transgenic Mouse Model of Cerebral Amyloid Angiopathy—A 9.4 Tesla MRI Study. Frontiers in Aging Neuroscience. 2016;8:1–9.

18. Improving Bioscience Research Reporting: The ARRIVE Guidelines for Reporting Animal Research | PLOS Biology [Internet]. [cited 2023 Dec 14];Available from: https://journals.plos.org/plosbiology/article?id=10.1371/journal.pbio.1000412

19. Munting LP, Derieppe MPP, Suidgeest E, Denis de Senneville B, Wells JA, van der Weerd L. Influence of different isoflurane anesthesia protocols on murine cerebral hemodynamics measured with pseudo-continuous arterial spin labeling. NMR Biomed. 2019;32:e4105.

20. Brossard C, Montigon O, Boux F, Delphin A, Christen T, Barbier EL, Lemasson B. MP3: Medical Software for Processing Multi-Parametric Images Pipelines. Front Neuroinform. 2020;14:594799.

21. Buxton RB, Frank LR, Wong EC, Siewert B, Warach S, Edelman RR. A general kinetic model for quantitative perfusion imaging with arterial spin labeling. Magn Reson Med. 1998;40:383–396.

22. Herscovitch P, Raichle ME. What is the correct value for the brain--blood partition coefficient for water? J Cereb Blood Flow Metab. 1985;5:65–69.

23. Dobre MC, Uğurbil K, Marjanska M. Determination of blood longitudinal relaxation time (T1) at high magnetic field strengths. Magn Reson Imaging. 2007;25:733–735.

24. Hirschler L, Collomb N, Voiron J, Köhler S, Barbier EL, Warnking JM. SAR comparison between CASL and pCASL at high magnetic field and evaluation of the benefit of a dedicated labeling coil. Magn Reson Med. 2020;83:254–261.

25. Sato Y, Nakajima S, Shiraga N, Atsumi H, Yoshida S, Koller T, Gerig G, Kikinis R. Three-dimensional multi-scale line filter for segmentation and visualization of curvilinear structures in medical images. Med Image Anal. 1998;2:143–168.

26. Ward JM, Vogel P, Sundberg JP. Brain and spinal cord lesions in 28 inbred strains of aging mice. Vet Pathol. 2022;59:1047–1055.

27. Iadecola C. Neurovascular regulation in the normal brain and in Alzheimer’s disease. Nature reviews Neuroscience. 2004;5:347–60.

28. Koemans EA, Chhatwal JP, van Veluw SJ, van Etten ES, van Osch MJP, van Walderveen MAA, Sohrabi HR, Kozberg MG, Shirzadi Z, Terwindt GM, van Buchem MA, Smith EE, Werring DJ, Martins RN, Wermer MJH, Greenberg SM. Progression of cerebral amyloid angiopathy: a pathophysiological framework. Lancet neurology. 2023;22:632–642.

29. Shin HK, Jones PB, Garcia-Alloza M, Borrelli L, Greenberg SM, Bacskai BJ, Frosch MP, Hyman BT, Moskowitz MA, Ayata C. Age-dependent cerebrovascular dysfunction in a transgenic mouse model of cerebral amyloid angiopathy. Brain. 2007;130:2310–2319.

30. Kim SH, Ahn JH, Yang H, Lee P, Koh GY, Jeong Y. Cerebral amyloid angiopathy aggravates perivascular clearance impairment in an Alzheimer’s disease mouse model. Acta Neuropathol Commun. 2020;8:181.

31. Schnerr RS, Jansen JFA, Uludag K, Hofman PAM, Wildberger JE, van Oostenbrugge RJ, Backes WH. Pulsatility of Lenticulostriate Arteries Assessed by 7 Tesla Flow MRI—Measurement, Reproducibility, and Applicability to Aging Effect. Frontiers in Physiology [Internet]. 2017 [cited 2023 Dec 14];8. Available from: https://www.frontiersin.org/articles/10.3389/fphys.2017.00961

32. Mitchell GF, van Buchem MA, Sigurdsson S, Gotal JD, Jonsdottir MK, Kjartansson Ó, Garcia M, Aspelund T, Harris TB, Gudnason V, Launer LJ. Arterial stiffness, pressure and flow pulsatility and brain structure and function: the Age, Gene/Environment Susceptibility – Reykjavik Study. Brain. 2011;134:3398–3407.

33. Takamura T, Motosugi U, Sasaki Y, Kakegawa T, Sato K, Glaser KJ, Ehman RL, Onishi H. Influence of Age on Global and Regional Brain Stiffness in Young and Middle-Aged Adults. J Magn Reson Imaging. 2020;51:727–733.

34. Mestre H, Tithof J, Du T, Song W, Peng W, Sweeney AM, Olveda G, Thomas JH, Nedergaard M, Kelley DH. Flow of cerebrospinal fluid is driven by arterial pulsations and is reduced in hypertension. Nature communications. 2018;9:4878–4878.

35. Maier FC, Wehrl HF, Schmid AM, Mannheim JG, Wiehr S, Lerdkrai C, Calaminus C, Stahlschmidt A, Ye L, Burnet M, Stiller D, Sabri O, Reischl G, Staufenbiel M, Garaschuk O, Jucker M, Pichler BJ. Longitudinal PET-MRI reveals β-amyloid deposition and rCBF dynamics and connects vascular amyloidosis to quantitative loss of perfusion. Nat Med. 2014;20:1485–1492.

36. Beckmann N, Gérard C, Abramowski D, Cannet C, Staufenbiel M. Noninvasive Magnetic Resonance Imaging Detection of Cerebral Amyloid Angiopathy-Related Microvascular Alterations Using Superparamagnetic Iron Oxide Particles in APP Transgenic Mouse Models of Alzheimer’s Disease: Application to Passive Aβ Immunotherapy. J Neurosci. 2011;31:1023–1031.

37. Kimchi EY, Kajdasz S, Bacskai BJ, Hyman BT. Analysis of Cerebral Amyloid Angiopathy in a Transgenic Mouse Model of Alzheimer Disease Using In Vivo Multiphoton Microscopy. Journal of Neuropathology & Experimental Neurology. 2001;60:274–279.

38. Dietrich HH, Horiuchi T, Xiang C, Hongo K, Falck JR, Dacey RG. Mechanism of ATP-induced local and conducted vasomotor responses in isolated rat cerebral penetrating arterioles. Journal of vascular research. 2009;46:253–64.

39. Nortley R, Korte N, Izquierdo P, Hirunpattarasilp C, Mishra A, Jaunmuktane Z, Kyrargyri V, Pfeiffer T, Khennouf L, Madry C, Gong H, Richard-Loendt A, Huang W, Saito T, Saido TC, Brandner S, Sethi H, Attwell D. Amyloid β oligomers constrict human capillaries in Alzheimer’s disease via signaling to pericytes. Science. 2019;365:eaav9518.

40. Meyer EP, Ulmann-Schuler A, Staufenbiel M, Krucker T. Altered morphology and 3D architecture of brain vasculature in a mouse model for Alzheimer’s disease. Proceedings of the National Academy of Sciences. 2008;105:3587–3592.

41. Chung Y-A, O JH, Kim J-Y, Kim K-J, Ahn K-J. Hypoperfusion and ischemia in cerebral amyloid angiopathy documented by 99mTc-ECD brain perfusion SPECT. J Nucl Med. 2009;50:1969–1974.

42. Gregg NM, Kim AE, Gurol ME, Lopez OL, Aizenstein HJ, Price JC, Mathis CA, James JA, Snitz BE, Cohen AD, Kamboh MI, Minhas D, Weissfeld LA, Tamburo EL, Klunk WE. Incidental Cerebral Microbleeds and Cerebral Blood Flow in Elderly Individuals. JAMA Neurology. 2015;72:1021–1028.

43. Eckrich J, Frenis K, Rodriguez-Blanco G, Ruan Y, Jiang S, Bayo Jimenez MT, Kuntic M, Oelze M, Hahad O, Li H, Gericke A, Steven S, Strieth S, von Kriegsheim A, Münzel T, Ernst BP, Daiber A. Aircraft noise exposure drives the activation of white blood cells and induces microvascular dysfunction in mice. Redox Biol. 2021;46:102063.

44. Frenis K, Kalinovic S, Ernst BP, Kvandova M, Al Zuabi A, Kuntic M, Oelze M, Stamm P, Bayo Jimenez MT, Kij A, Keppeler K, Klein V, Strohm L, Ubbens H, Daub S, Hahad O, Kröller-Schön S, Schmeisser MJ, Chlopicki S, Eckrich J, Strieth S, Daiber A, Steven S, Münzel T. Long-Term Effects of Aircraft Noise Exposure on Vascular Oxidative Stress, Endothelial Function and Blood Pressure: No Evidence for Adaptation or Tolerance Development. Front Mol Biosci. 2022;8:814921.

45. Marazuela P, Paez-montserrat B, Bonaterra-pastra A, Hern M, Sol M. Impact of Cerebral Amyloid Angiopathy in Two Transgenic Mouse Models of Cerebral β -Amyloidosis : A Neuropathological Study. 2022;

46. Eyre B, Shaw K, Sharp P, Boorman L, Lee L, Shabir O, Berwick J, Howarth C. The effects of locomotion on sensory-evoked haemodynamic responses in the cortex of awake mice. Sci Rep. 2022;12:6236.

47. Huo B-X, Smith JB, Drew PJ. Neurovascular coupling and decoupling in the cortex during voluntary locomotion. J Neurosci. 2014;34:10975–10981.

48. Turner KL, Gheres KW, Proctor EA, Drew PJ. Neurovascular coupling and bilateral connectivity during NREM and REM sleep. eLife. 2020;9:e62071.

