## Supplemental Material for "Loss of spontaneous vasomotion precedes impaired cerebrovascular reactivity and microbleeds in a mouse model of cerebral amyloid angiopathy"

**Supplemental Material - Detailed Methods**

Mouse housing

Mice in both the 2-photon and MRI cohorts were housed together (2 – 4 littermates per cage), except for the mice in 2-photon cohorts after they underwent craniotomy. They were then housed individually to prevent damaging of the cranial implants by cage mates. Housing consisted of individually ventilated cages, supplied with bedding, an igloo, and unlimited chow food and water.

*2-photon microscopy*

Longitudinal imaging cohort:
- 4 WT mice (2 bred in-house, 1 female)
- 4 Tg mice (2 bred in-house, 2 female)

15–16-month-old cohort:
- 4 WT (0 bred in-house, 1 female)
- 3 Tg (0 bred in-house, 2 female)

Isoflurane titration cohort (15-16-month-old):
- 3 WT (3 bred-in house, 1 female)
- 3 Tg (3 bred in-house, 1 female)

*MRI*

Cross-sectional:
- 12 months: 6 WT mice (0 bred in-house, 4 female), 5 Tg mice (0 bred in-house, 4 female)
- 18 months: 10 WT mice (4 bred in-house, 5 female), 10 Tg mice (5 bred in-house, 4 female)
- 24 months: 9 WT littermates (8 bred in-house, 3 female), 6 Tg (5 bred in-house, 4 female)

*Immunohistochemistry*

Cross-sectional:
- 12 months: 3 WT (3 bred in-house, 2 female), 3 Tg (3 bred in-house, 3 female)
- 18 months: 7 WT (4 bred in-house, 3 female), 6 Tg (4 bred in-house, 3 female)
- 24 months: 8 WT (8 bred in-house, 3 female), 5 Tg (5 bred in-house, 3 female)

Note, all 12-month-old mice (both WT and Tg) had cranial windows placed during life (1 WT and 2 Tg mice imaged as a part of the longitudinal 2-photon imaging cohort described above, and the others with smaller windows placed for a separate set of longitudinal imaging experiments). None of the 18- and 24-month-old mice had cranial windows placed, as these mice were all imaged in the MRI experiments.

Image acquisition – 2-photon

The same 4 fields of view (FOVs) were imaged in each session, unless clouding of the cranial window prevented clear visualization of the vasculature (3-6 imaging sessions/mouse). In each imaging session, Z-stacks to assess vascular structure and vascular amyloid-β coverage were acquired at a speed of 4 μs/pixel, a pixel resolution of 1 $\mu m^2$ (512x512 pixels), and a step size of 5 μm at 1x magnification. Spontaneous vasomotion was recorded in pial arterioles continuously over ~ 5 min at a speed of 2 μs/pixel, pixel resolution of 2 $\mu m^2$ (256x256 pixels) and a frame rate of ~2.3 Hz at 1x magnification. Four FOVs were obtained per mouse (2 in each hemisphere). Line scans were performed in arteriolar and venular segments in the same FOV using bidirectional scanning at a speed of 0.244 ms/line at 6x magnification for 10,000 lines, both perpendicular to the vessel wall (to assess diameter) and parallel to the vessel wall (to assess red blood cell [RBC] velocity). PMT settings were the same throughout all imaging sessions, and laser power was adjusted as needed.

Image acquisition – MRI

The total scan duration was approximately 1.5 hours. First, 1-minute anatomical $T_2$-weighted Rapid Acquisition with Relaxation Enhancement (RARE) scans were acquired in all 3 directions for consistent planning of the subsequent sequences. pCASL label and control inter-pulse phases were optimized with pre-scans[1]. Thereafter, a 15-minute pCASL scan was acquired, with the following parameters: repetition time (TR) / echo time (TE) = 3,500 / 28 ms, labeling duration ($\tau$) = 2,950 ms, post-labeling delay (PLD) = 300 ms, 5 slice spin-echo echo-planar imaging (EPI) read-out, 0.25 x 0.25 x 1.0 $mm^3$ resolution and a 0.5 mm slice gap. During the 15-minute pCASL scan, the vasculature was challenged by adding 10 % $CO_2$ to the isoflurane gas mixture from minute 5 to 10. Oxygen and isoflurane concentrations were kept constant during the challenge. Two additional scans were acquired to support cerebral blood flow (CBF) quantification: an inversion recovery sequence with the same EPI read-out parameters as the pCASL was acquired to estimate tissue $T_1$ ($T_{1t}$) and tissue magnetization ($M_{0t}$), and a pCASL flow-compensated, fast low angle shot (fc-FLASH) was acquired at the level of the carotids, 3 mm downstream of the labeling plane, to determine labeling efficiency ($\alpha$). An additional $T_2$-weighted RARE scan with the same geometry as the pCASL scan was acquired for registration purposes. To visualize the large arteries, a 3D time-of-flight (TOF) scan was acquired, covering the head and neck region at a 0.10 x 0.10 x 0.16 $mm^3$ resolution. Lastly, to detect microbleeds, an MGE sequence was acquired, which consisted of 6 echoes (3.5 – 21 msec, with 3.5 msec intervals), a resolution of 0.175 x 0.175 x 0.35 $mm^3$ and 25 consecutive slices.

Image processing – MRI

For CBF quantification, the labeling efficiency ($\alpha$) was assumed to be 0.80, which was the median $\alpha$ of all WT animals (Supplemental Figure 6) and is in correspondence with literature[2]. We chose not to use each mouse's individual $\alpha$ value – derived by calculating the relative complex signal difference between label and control images[1] - in the CBF quantification, to prevent introduction of extra noise in the CBF estimation. The measurement was noisy because the carotids were located too deep to be imaged with the surface coil. The pCASL fc-FLASH sequence was therefore carried out with the whole-body volume transmit coil, which has a lower signal-to-noise ratio. The labeling efficiency ($\alpha$) could therefore only be determined in a couple of carotid voxels. Nevertheless, we used the measurements to assess if there were large differences in $\alpha$ on the group level between the genotypes, which was not the case (Supplemental Figure 6).

To put all CBF and CVR maps in the same space, the aligned pCASL EPIs of each mouse were first registered to its corresponding $T_2$-weighted RARE scan with the same geometry, and then to an arbitrarily chosen reference $T_2$-weighted RARE of 1 WT mouse, using Evolution software[3]. This process has been described in detail before[4].

Immunohistochemistry

Sections were deparaffinized and rehydrated through xylene and graded series of ethanol (100 %, 95 %, 70 %) and tap water. Then, sections were treated with 3 % $H_2O_2$ to quench endogenous peroxidase activity. For SMA, antigen retrieval was performed with heat induced epitope retrieval in citrate buffer (pH 6.0). For amyloid-β, either 100 % formic acid (5 min) or heated citrate buffer was used for antigen retrieval (20 min). The sections were blocked using either normal goat serum (SMA) or normal horse serum (amyloid-β) diluted in tris buffered saline (TBS) from a Vectastain ABC kit (Vector Laboratories, cat.nr. # PK-4002) for 1 hour. Next, sections were incubated overnight in a 4°C cold room with primary antibodies against SMA (mouse monoclonal, Dako, cat.nr. # M0851, 1:250) or amyloid-β (mouse monoclonal, Dako, cat.nr. # M0872, 1:200) diluted in TBS. The following day, a biotinylated secondary antibody was applied from a Vectastain ABC kit (Vector Laboratories). An avidin-biotin complex was applied, and the substrate was visualized using the chromogen 3,3'-Diaminobenzidine (DAB) (Vector Laboratories, cat.nr. # SK-4100). The sections were counterstained with haematoxylin. The sections were washed using TBS before and after every secondary antibody incubation and after avidin-biotin application. Sections were then dehydrated through a series of ethanol (70 %, 95 %, 100 %), cleared through xylene, and cover slipped using Permount mounting medium (Fisher Chemical, cat.nr. # SP15-100).

Statistics

To compare baseline arteriolar diameter, arteriolar RBC velocity, and heart rate, student's t-tests were used in Prism v10. Each vessel segment was treated as independent for this analysis. To compare the maximum amplitude of the vasomotion peak between Tg and WT mice across 3 age groups (7-9 months, 12-14 months, and 15-16 months), a one-way ANOVA test was performed in Prism v10. Using a Šídák correction, post-hoc multiple comparisons were performed on preselected pairs as follows: 7-9mo-WT vs 7-9mo-Tg, 12-14mo-WT vs 12-14mo-Tg, and 15-16mo-WT vs 15-16mo-Tg. To compare the peak vasomotion frequency between WT and Tg mice at the first (7-9 months old) and last (12-14 months old) imaging sessions, student's t-tests in Prism v10 were used (the average vasomotion frequency for each mouse imaging session was used for this analysis).

To compare CBF and CVR between the two genotypes, Mann-Whitney U tests were performed using MATLAB's "ranksum" function. The mean CBF and CVR values per mouse were used, which were derived respectively by taking the average CBF between minute 1-5, and the percentage CBF increase in minute 6-10 over minute 1-5. Spearman's rank-order correlations were performed for the relationship between microbleed count and CBF/CVR in Tg mice.

To compare vascular SMA coverage across groups, a one-way ANOVA was performed in Prism v10. Using a Šídák correction, post-hoc multiple comparisons were performed on preselected pairs as follows: 12mo-WT vs 12mo-Tg, 18mo-WT vs 18mo-Tg, 24mo-WT vs 24mo-Tg, 12mo-WT vs 18mo-WT, 12mo-WT vs 24mo-WT, 18mo-WT vs 24mo-WT, 12mo-Tg vs 18mo-Tg, 12mo-Tg vs 24mo-Tg, 18mo-Tg vs 24mo-Tg.

Supplemental Figures

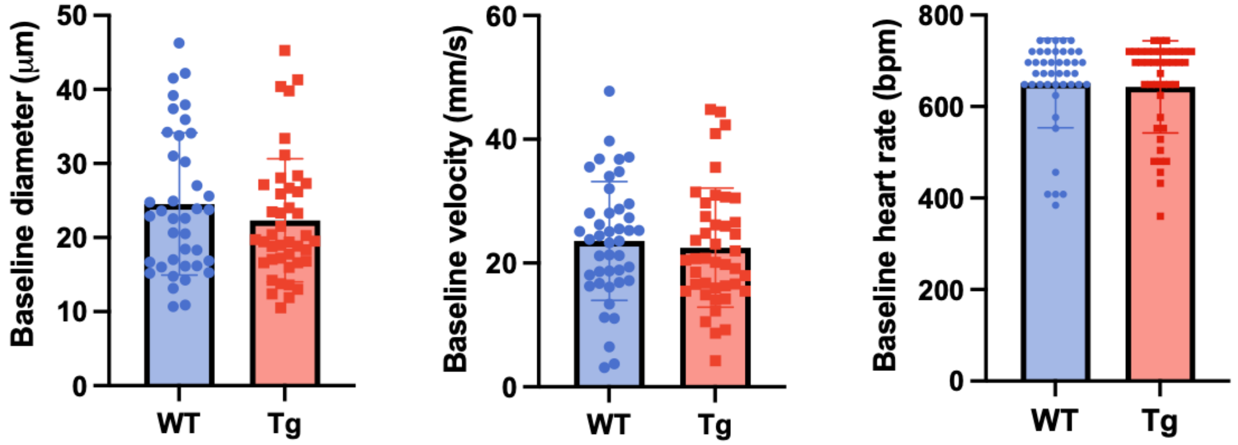

**Supplemental Figure 1. No difference in baseline arteriolar diameter, velocity, or heart rate between APP23 Tg mice and WT littermates.** Mean +/- SD shown. Left: n = 39, 42 arterioles; middle: n = 43, 42 arterioles; right: n = 43, 42 arterioles (note, each measurement treated as independent).

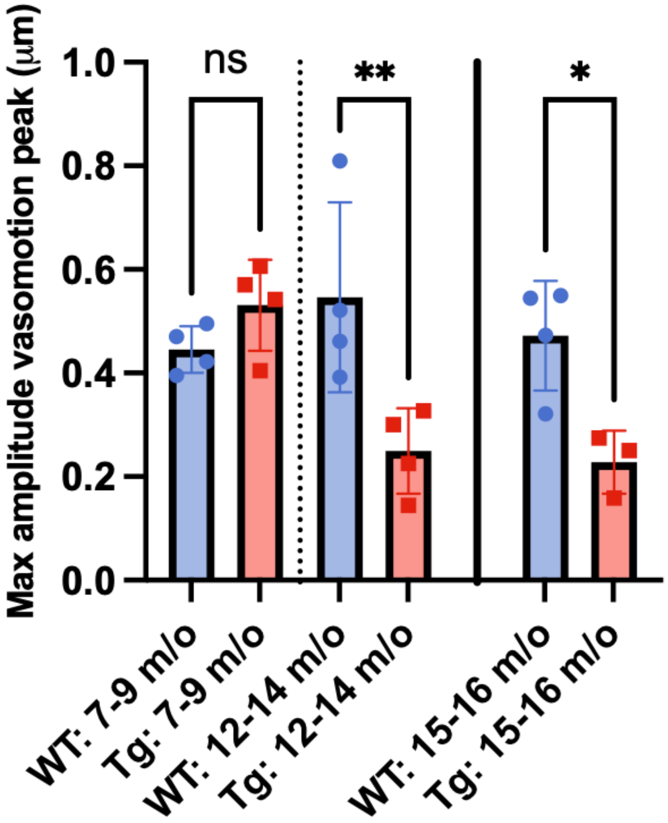

**Supplemental Figure 2. Vasomotion remains intact in a separate group of WT mice at 15-16 months of age.** The same cohort of mice was imaged between 7-14 months (n = 4 WT, 4 Tg). The 15–16-month-old mice shown represent an independent cohort of mice (n = 4 WT, 3 Tg). One-way ANOVA ($p = 0.0018$), pre-selected comparisons performed using Šídák correction shown (* $p < 0.05$, ** $p < 0.01$).

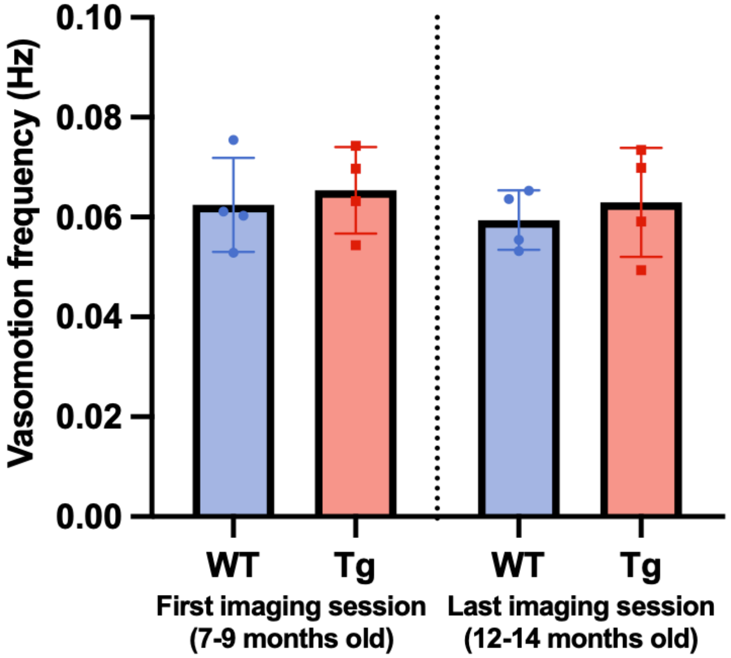

**Supplemental Figure 3. No difference in vasomotion frequency between APP23 Tg and WT mice.** Frequency does not change over the imaged age range (n = 4 WT, 4 Tg mice).

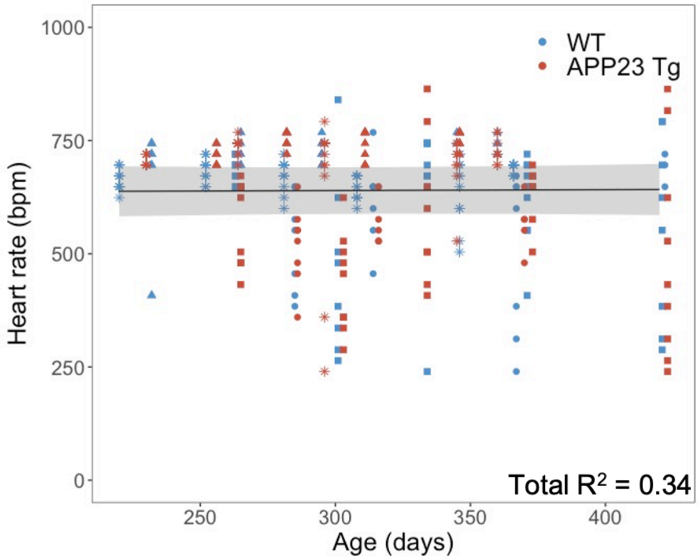

**Supplemental Figure 4. No change in heart rate with age in APP23 Tg and WT mice.** Each mouse is depicted with a different symbol (n = 4 Tg, 4 WT mice, 358 measurements across mice). Line represents LME model including mouse and vessel as random effects and age as a fixed effect (shaded areas represent the 95% confidence interval of the model's prediction, Model 1, see Supplemental Table 6). No significant association with age was observed in this model.

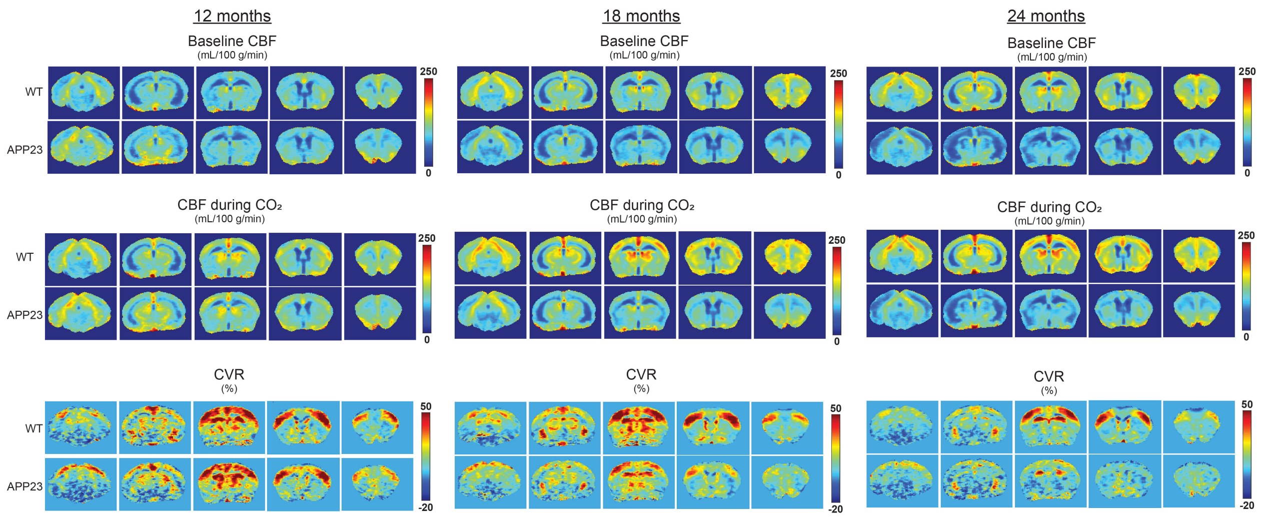

**Supplemental Figure 5. Whole-brain MRI maps of cerebral blood flow (CBF) and cerebrovascular reactivity (CVR).** The rows display maps averaged per genotype (wild type [WT] or APP23 transgenic [Tg]) and per age group (12, 18 or 24 months old), with each group consisting of n = 5-9 mice. On the columns, the maps display the baseline CBF, CBF during $CO_2$, and CVR (relative CBF change during $CO_2$), and in the sub-columns the different 5 MRI slices are shown.

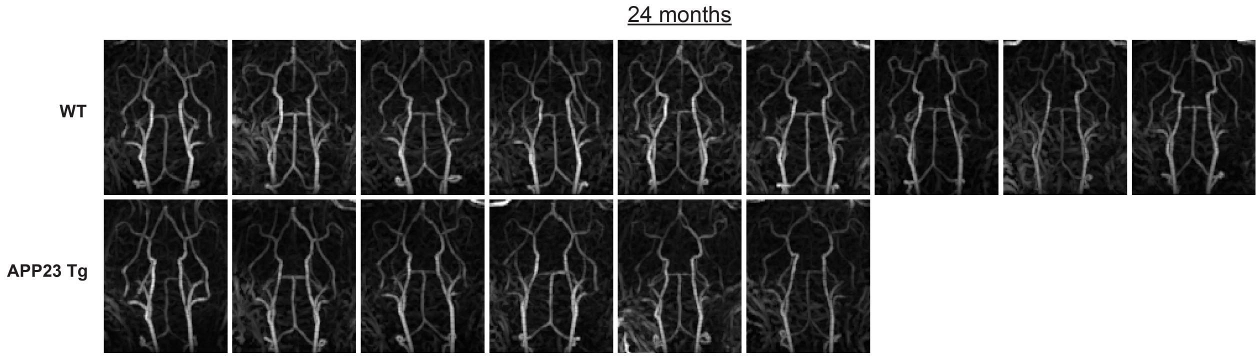

**Supplemental Figure 6. MRI time-of-flight (TOF) images of all mice in the 24-month-old cohort.** Maximum intensity projections of vessel enhancement filtered TOF images are shown for wild type (WT) mice on the top row, and for APP23 transgenic (Tg) mice on the bottom row.

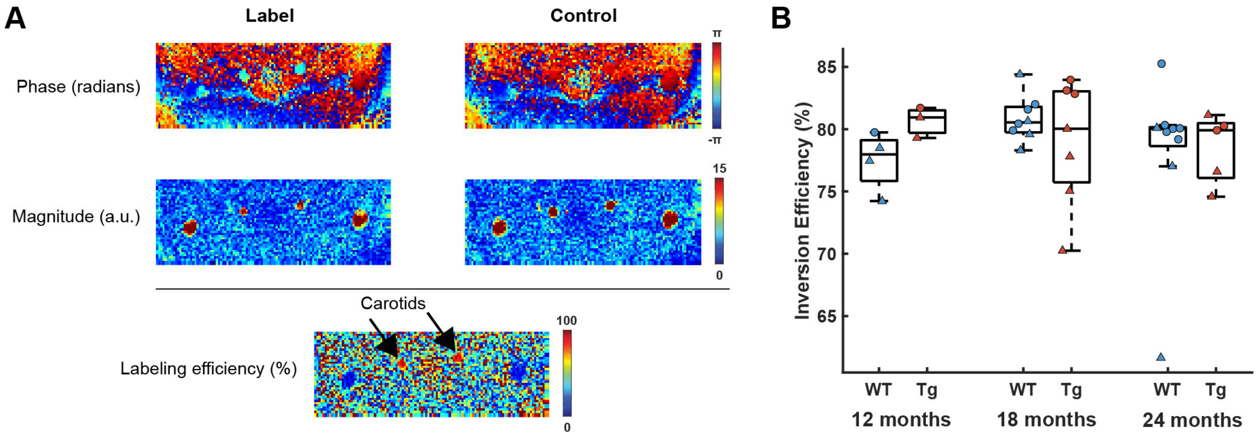

**Supplemental Figure 7. Pseudo-continuous arterial spin labeling (pCASL) inversion efficiency measurements.** Phase and magnitude images of the pCASL fc-FLASH sequence acquired at the level of the carotids are shown in A), both for label and control acquisitions. The relative complex signal difference between label and control images is shown on the bottom image in A), where the right and left carotids are indicated with black arrows. The plot in B) displays the inversion efficiency values measured in the carotids for the 12, 18, and 24-month-old cohorts, showing no significant differences between wild type (WT) and APP23 transgenic (Tg) mice. Dots represent male and triangles represent female mice.

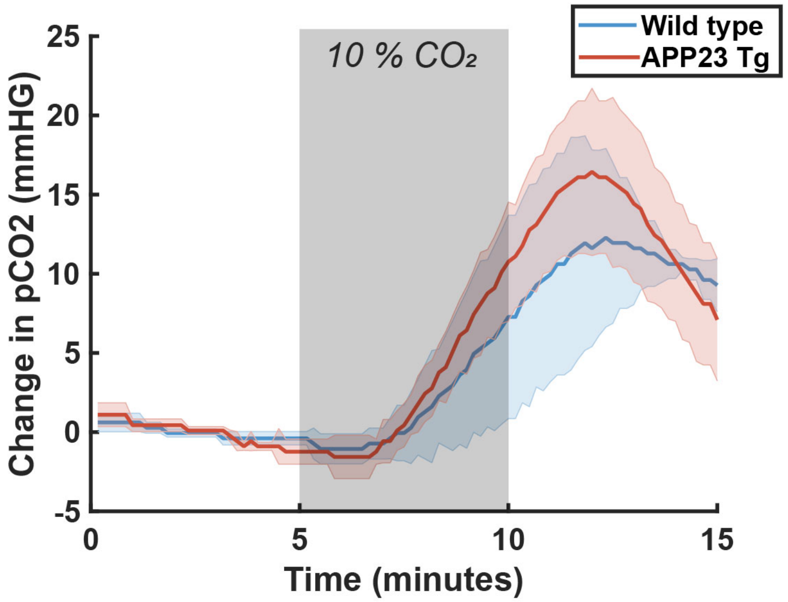

**Supplemental Figure 8. Transcutaneous $pCO_2$ (tc-$pCO_2$) measurements.** The graphs show the mean mmHg change in tc-pCO2 (± standard deviation) during a 10% CO2 challenge, acquired in 3 wild type and 3 APP23 transgenic (Tg) mice from the 18-month-old cohort, which underwent a separate 10% $CO_2$ challenge outside the MRI scanner.

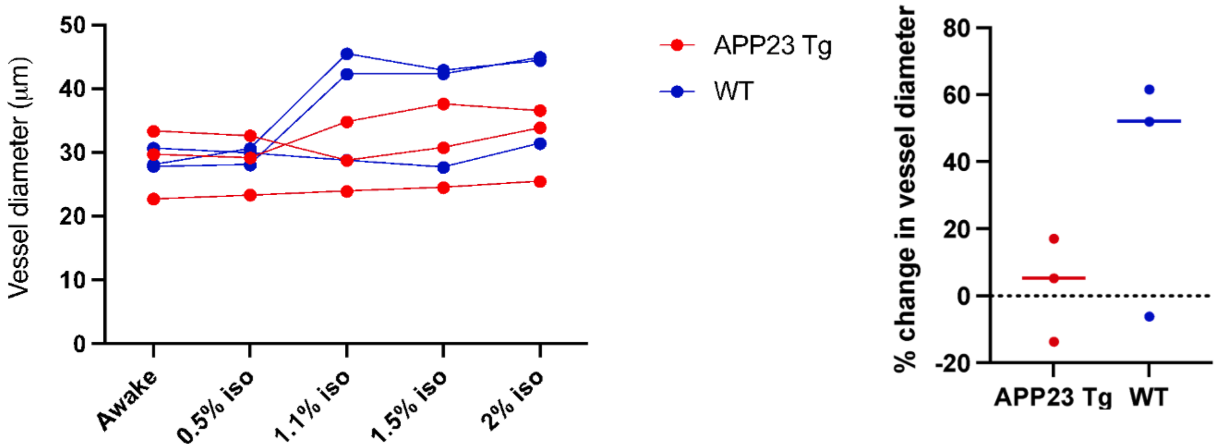

**Supplemental Figure 9. Baseline arteriolar diameter increases more strongly with isoflurane in 15-month-old WT compared to APP23 Tg mice.** Left: measurements of same arterioles imaged in APP23 Tg and WT mice with increasing isoflurane administered. Right: % change in vessel diameter from awake imaging to 1.1% isoflurane in APP23 Tg and WT mice.

**Supplemental Tables**

**Supplemental Table 1. CAA coverage (%) model**

| | | Null model | **Model 1** |
|---|---|---|---|
| Age | p-value<br>estimate<br>95% CI | | **p = 9.7E-16**<br>4.60E-01<br>[0.38 0.54] |
| Log likelihood<br>AIC<br>BIC<br>Total R2 | | -298.3<br>604.5<br>613<br>0.34 | **-265.4**<br>**540.8**<br>**551.4**<br>**0.8** |

Results for the fixed effects estimates of LME models developed with % CAA coverage as the dependent variable. In all models, mouse and vessel were treated as random effects. Model 1, including age as a fixed effect, had the best fit (n = 62 measurements across 4 Tg mice).

**Supplemental Table 2. Normalized arteriolar diameter models**

| | | Null model | Model 1 | Model 2 | **Model 3** |
|---|---|---|---|---|---|
| Age | p-value<br>estimate<br>95% CI | | **p = 0.013**<br>2.0E-04<br>[4.1E-5 3.6E-3] | **p = 0.017**<br>1.9E-04<br>[3.1E-5 3.5E-4] | p = 0.94<br>-7.1E-06<br>[-2.1E-4 2E-4] |
| Genotype (Tg) | p-value<br>estimate<br>95% CI | | | **p = 0.046**<br>3.3E-02<br>[2.7E-3 6.5E-2] | **p = 0.037**<br>-1.0E-01<br>[-2E-1 -2.4E-3] |
| Age*Genotype (Tg) | p-value<br>estimate<br>95% CI | | | | **p = 0.0051**<br>4.4E-04<br>[1.2E-4 7.5E-4] |
| Log likelihood<br>AIC<br>BIC<br>Total R2 | | 390.2<br>-772.4<br>-756.9<br>0.23 | 393.2<br>-776.4<br>-757.1<br>0.25 | 395.4<br>-778.8<br>-755.6<br>0.25 | **399.1**<br>**-784.2**<br>**-757.1**<br>**0.26** |

Results for the fixed effects estimates of LME models developed with normalized arteriolar diameter as the dependent variable. In all models, mouse and vessel were treated as random effects. Model 3, including age, genotype, and age*genotype interactions as fixed effects had the best fit (n = 355 vessel measurements across 4 Tg and 4 WT mice).

**Supplemental Table 3. Normalized RBC velocity models**

| | | **Null model** | Model 1 | Model 2 | Model 3 |
|---|---|---|---|---|---|
| Age | p-value | | p = 0.68 | p = 0.65 | p = 0.66 |
| | estimate | | 5.5E-05 | 6.0E-05 | -7.5E-06 |
| | 95% CI | | [-2.2E-4 3.2E-4] | [-2.0E-3 3.2E-4] | [-4.1E-4 2.6E-4] |
| Genotype (Tg) | p-value | | | p = 0.088 | p = 0.41 |
| | estimate | | | 3.7E-02 | -7.1E-02 |
| | 95% CI | | | [-6.9E-3 7.9E-2] | [-2.4E-1 9.9E-2] |
| Age*Genotype (Tg) | p-value | | | | p = 0.2 |
| | estimate | | | | 3.5E-04 |
| | 95% CI | | | | [-1.8E-4 8.9E-4] |
| Log likelihood | | **187.1** | 187.2 | 188.7 | 189.5 |
| AIC | | **-366.3** | -364.4 | -365.3 | -365 |
| BIC | | **-351** | -345.4 | -342.4 | -338.3 |
| Total R2 | | **0.28** | 0.28 | 0.28 | 0.27 |

Results for the fixed effects estimates of LME models developed with normalized RBC velocity as the dependent variable. In all models, mouse and vessel were treated as random effects. The null model (no fixed effects) had the best fit (n = 334 vessel measurements across 4 Tg and 4 WT mice).

**Supplemental Table 4. Pulsatility (area under curve) models**

| | | Null model | Model 1 | Model 2 | Model 3 | **Model 4** |
|---|---|---|---|---|---|---|
| Diameter | p-value | | **p = 7.51E-12** | **p = 4.39E-10** | **p = 4.6E-10** | **p = 3.8E-9** |
| | estimate | | 8.50E-03 | 7.73E-03 | 7.71E-03 | 7.33E-03 |
| | 95% CI | | [0.0061 0.011] | [0.0055 0.010] | [0.0055 0.010] | [0.0051 0.0096] |
| Age | p-value | | | **p = 2.9E-11** | **p = 2.3E-11** | **p = 0.0012** |
| | estimate | | | 1.59E-03 | 1.60E-03 | 9.73E-04 |
| | 95% CI | | | [0.0011 0.0020] | [0.0011 0.0020] | [0.00039 0.0016] |
| Genotype (Tg) | p-value | | | | p = 0.327 | **p = 0.040** |
| | estimate | | | | 1.06E-01 | 3.60E-01 |
| | 95% CI | | | | [-0.12 0.33] | [-0.71 - 0.014] |
| Age*Genotype (Tg) | p-value | | | | | **p = 0.0012** |
| | estimate | | | | | 1.50E-03 |
| | 95% CI | | | | | [0.00059 0.0024] |
| Log likelihood | | -8.7 | 14.7 | 37 | 37.5 | **42.7** |
| AIC | | 25.3 | -19.1 | -62 | -61 | **-69.4** |
| BIC | | 40.8 | -0.1 | -38.7 | -33.9 | **-38.4** |
| Total R2 | | 0.37 | 0.43 | 0.46 | 0.46 | **0.48** |

Results for the fixed effects estimates of LME models developed with pulsatility (measured as area under the curve) as the dependent variable. In all models, mouse and vessel were treated as random effects. Model 4, including vessel diameter, age, genotype, and age*genotype interactions as fixed effects, had the best fit (n = 355 vessel measurements across 4 Tg and 4 WT mice).

**Supplemental Table 5. Pulsatility (peak amplitude) models**

| | | Null model | Model 1 | **Model 2** | Model 3 | Model 4 |
|---|---|---|---|---|---|---|
| Diameter | p-value | | **p = 2.8E-14** | **p = 4.0E-13** | **p = 4.5E-13** | **p = 6.8E-13** |
| | estimate | | 1.30E-02 | 1.25E-02 | 1.24E-02 | 1.25E-02 |
| | 95% CI | | [0.010 0.016] | [0.0094 0.015] | [0.0094 0.015] | [0.0095 0.016] |
| Age | p-value | | | **p = 0.0077** | **p = 0.0068** | **p = 0.015** |
| | estimate | | | 7.08E-04 | 7.21E-04 | 7.96E-04 |
| | 95% CI | | | [0.00019 0.0012] | [0.00019 0.0013] | [0.00016 0.0014] |
| Genotype (Tg) | p-value | | | | p = 0.57 | p = 0.61 |
| | estimate | | | | 1.80E-02 | 8.44E-02 |
| | 95% CI | | | | [-0.046 0.089] | [-0.25 0.41] |
| Age*Genotype (Tg) | p-value | | | | | p = 0.69 |
| | estimate | | | | | -2.21E-04 |
| | 95% CI | | | | | [-0.0013 0.00086] |
| Log likelihood | | 9 | 37.4 | **40.8** | 41 | 41 |
| AIC | | -10.1 | -64.5 | **-69.6** | -67.9 | -66.1 |
| BIC | | 3.9 | -47.1 | **-48.7** | -43.5 | -38.1 |
| Total R2 | | 0.41 | 0.44 | **0.43** | 0.43 | 0.44 |

Results for the fixed effects estimates of LME models developed with pulsatility (measured as peak amplitude at the heart rate frequency in the frequency domain) as the dependent variable. In all models, mouse and vessel were treated as random effects. Model 2, including vessel diameter and age as fixed effects, had the best fit (n = 242 measurements across 4 Tg and 4 WT mice).

**Supplemental Table 6. Heart rate models**

| | | **Null model** | Model 1 | Model 2 | Model 3 |
|---|---|---|---|---|---|
| Age | p-value | | p = 0.86 | p = 0.86 | p = 0.999 |
| | estimate | | 2.00E-02 | 2.00E-02 | 2.10E-04 |
| | 95% CI | | [-0.20 0.24] | [-0.20 0.24] | [-0.29 0.29] |
| Genotype (Tg) | p-value | | | p = 0.999 | p = 0.87 |
| | estimate | | | 3.66E-02 | -1.50E+01 |
| | 95% CI | | | [-116.2 115.8] | [-190.2 159.8] |
| Age*Genotype (Tg) | p-value | | | | p = 0.84 |
| | estimate | | | | 4.70E-02 |
| | 95% CI | | | | [-0.40 0.49] |
| Log likelihood | | **-2171.9** | -2171.9 | -2171.9 | -2171.9 |
| AIC | | **4351.9** | 4353.8 | 4355.8 | 4357.8 |
| BIC | | **4367.4** | 4373.2 | 4379.1 | 4384.9 |
| Total R2 | | **0.33** | 0.34 | 0.34 | 0.34 |

Results for the fixed effects estimates of LME models developed with heart rate as the dependent variable. In all models, mouse and vessel were treated as random effects. The null model (including no fixed effects) had the best fit (n = 242 measurements across 4 Tg and 4 WT mice).

**Supplemental Material - References**

1. Hirschler L, Debacker CS, Voiron J, Köhler S, Warnking JM, Barbier EL. Interpulse phase corrections for unbalanced pseudo-continuous arterial spin labeling at high magnetic field. *Magn Reson Med*. 2018;79:1314–1324.

2. Hirschler L, Collomb N, Voiron J, Köhler S, Barbier EL, Warnking JM. SAR comparison between CASL and pCASL at high magnetic field and evaluation of the benefit of a dedicated labeling coil. *Magn Reson Med*. 2020;83:254–261.

3. Denis de Senneville B, Zachiu C, Ries M, Moonen C. EVolution: an edge-based variational method for non-rigid multi-modal image registration. *Phys Med Biol*. 2016;61:7377–7396.

4. Munting LP, Derieppe MPP, Suidgeest E, Denis de Senneville B, Wells JA, van der Weerd L. Influence of different isoflurane anesthesia protocols on murine cerebral hemodynamics measured with pseudo-continuous arterial spin labeling. *NMR Biomed*. 2019;32:e4105.
